# Retracing the horizontal transfer of a novel innate immune factor in *Drosophila*

**DOI:** 10.1101/2024.05.29.596511

**Authors:** Rebecca L. Tarnopol, Josephine Tamsil, Gyöngyi Cinege, Ji Heon Ha, Kirsten I. Verster, Edit Ábrahám, Lilla B. Magyar, Bernard Y. Kim, Susan L. Bernstein, Zoltán Lipinszki, István Andó, Noah K. Whiteman

## Abstract

Immune systems are among the most dynamically evolving traits across the tree of life, and long-lived macroparasites play an outsized role in shaping animal immunity. Even without adaptive immunity, insects have evolved potent innate immune strategies to neutralize such enemies, including nematodes and parasitoid wasps. One such strategy relies on endosymbioses between insects and toxin-expressing bacteria. Here, we use genome editing in *Drosophila melanogaster* to retrace the evolution of two of such toxins — *cytolethal distending toxin B* (*cdtB*) and *apoptosis inducing protein of 56kDa* (*aip56*) — that were horizontally transferred from bacteriophages to insects. We found that a *cdtB::aip56* fusion gene (*fusionB*), which is conserved in *Drosophila ananassae* subgroup species, dramatically promoted fly survival and suppressed wasp development when expressed in *D. melanogaster* immune tissues. FusionB, a functional nuclease, was secreted into the host hemolymph where it targeted the parasitoid embryo’s serosal tissue and is to our knowledge the first humoral anti-parasitoid toxin in *Drosophila*. When expressed ubiquitously, *fusionB* slowed development in late stage fly larvae and eventually killed flies, pointing to the salience of regulatory constraint in preventing autoimmunity. Our findings demonstrate how horizontal gene transfer, in the right regulatory context, can instantly provide new and potent innate immune modules in animals.

## Introduction

Invertebrates and vertebrates share deeply homologous innate immune pathways effective against a suite of microbial pathogens^1–6^. In addition, invertebrates and vertebrates face attack by macroparasitic animals^7^ that cause chronic, long-term infections^8^, facilitating Red Queen-like host-parasite coevolutionary dynamics^7,9–11^. Indeed, the local community compositions of directly transmitted parasitic worms are the most important factor shaping genome-wide patterns of local adaptation in humans^12^. Many of these loci are also associated with autoimmunity^12^, indicating that antagonistic pleiotropy is a cost of “trench warfare”^13^ between hosts and macroparasites.

Among the most important macroparasites of insects are parasitoid wasps and nematodes. The parasitoid wasps and nematodes that infect *Drosophila* spp. larvae reach high prevalences in natural populations and can locally cause up to 75% mortality and 30% sterility, respectively^14,15^. Immune defenses against these enemies are rapidly evolving^16,17^ and costly^18^. As with parasitic worm infections in humans, *Drosophila* hosts deploy endogenous cellular and humoral responses against these parasitoids^19^, but also use externally acquired toxins through both defensive mutualisms with endosymbiotic bacteria expressing ribosome inactivating proteins (RIPs)^20,21^ and the dietary sequestration of ethanol^22,23^.

Cell-mediated immune responses to parasitoid wasps are well characterized in *Drosophila^24,25^.* Despite a wealth of information on humoral factors against microbes^6^, our understanding of their humoral responses against parasitoid wasps is far more limited. Cellular factors are associated with encapsulation and melanization of parasitoid neonates in *D. melanogaster*^24,26,27^, but toxic humoral factors against parasitoid wasps of *Drosophila* are unknown. However, putative toxic humoral factors have been identified in Lepidoptera that act against parasitoid wasps, including one acquired through horizontal gene transfer (HGT) from viruses^28,29^. More generally, HGT can instantaneously endow recipient lineages with new genes from donor lineages^30^ and has played an important role in the evolution of prokaryotic immune systems^31–33^. Yet, the role of HGT in the evolution of the animal immune system is less clear^34,35^.

We addressed this gap by leveraging an exceptional HGT event from ancestors of APSE phages or prophages of endosymbiotic bacteria to insects. Two horizontally transferred genes encoding eukaryotic toxins — *cytolethal distending toxin B* (*cdtB*) and *apoptosis inducing protein of 56kDa* (*aip56*) — were repeatedly transferred to the nuclear genomes of insects from five different orders^36–38^. *cdtB* encodes the catalytically active subunit of the tripartite cytolethal distending toxin (CDT) holotoxin^39^. CdtB is a DNAse I-type nuclease and causes G2/M cell cycle arrest in eukaryotic cells, eventually leading to apoptosis. AIP56 is a single polypeptide AB toxin that was first isolated from the fish pathogen *Photobacterium damselae*^40^. *P. damselae* AIP56 toxin is a metalloprotease that cleaves the eukaryotic NF-kB p65, resulting in cell death via apoptosis.

Both *cdtB* and *aip56* were horizontally transferred to a recent common ancestor of *Drosophila ananassae* subgroup fruit flies. The nuclear genome of *D. ananassae* encodes one full-length *cdtB* homolog adjacent to two *cdtB::aip56* “fusion” genes (henceforth *fusionA* and *fusionB,* or *fusA* and *fusB*), consisting of a full-length *cdtB* copy fused to the non-catalytic B domain of *aip56*^36,38^. The most closely related homologs are found in toxin cassette regions of APSE phage or prophage genomes that in turn infect or are encoded in the genomes of secondary bacterial endosymbionts of aphids and other sap-feeding insects. In these endosymbionts, *cdtB* and *aip56* provide hosts with protection against parasitoid wasp challenge (Figure 1A-C)^41–43^. Strikingly, *D. ananassae* has naturally high resistance to parasitoid wasp challenge compared to other drosophilids^16^, despite lacking blood cell types and a key prophenoloxidase gene (PPO3) that participate in these immune reactions in *D. melanogaster*^44^. Functional studies in *D. ananassae* demonstrated the involvement of horizontally transferred *cdtB* and *aip56* genes in parasitoid wasp defense. The genes are expressed primarily in the larval fat body, a key insect immune tissue, and are highly induced upon parasitoid attack^36,38^. Loss-of-function studies indicated these genes are necessary for *D. ananassae* to mount its full immune response against parasitoids (Figure 1D-E)^38^. Thus, this HGT event illuminates an important link between the evolution of defensive mutualisms that rely on toxins and humoral immunity in *Drosophila*.

**Figure 1.**
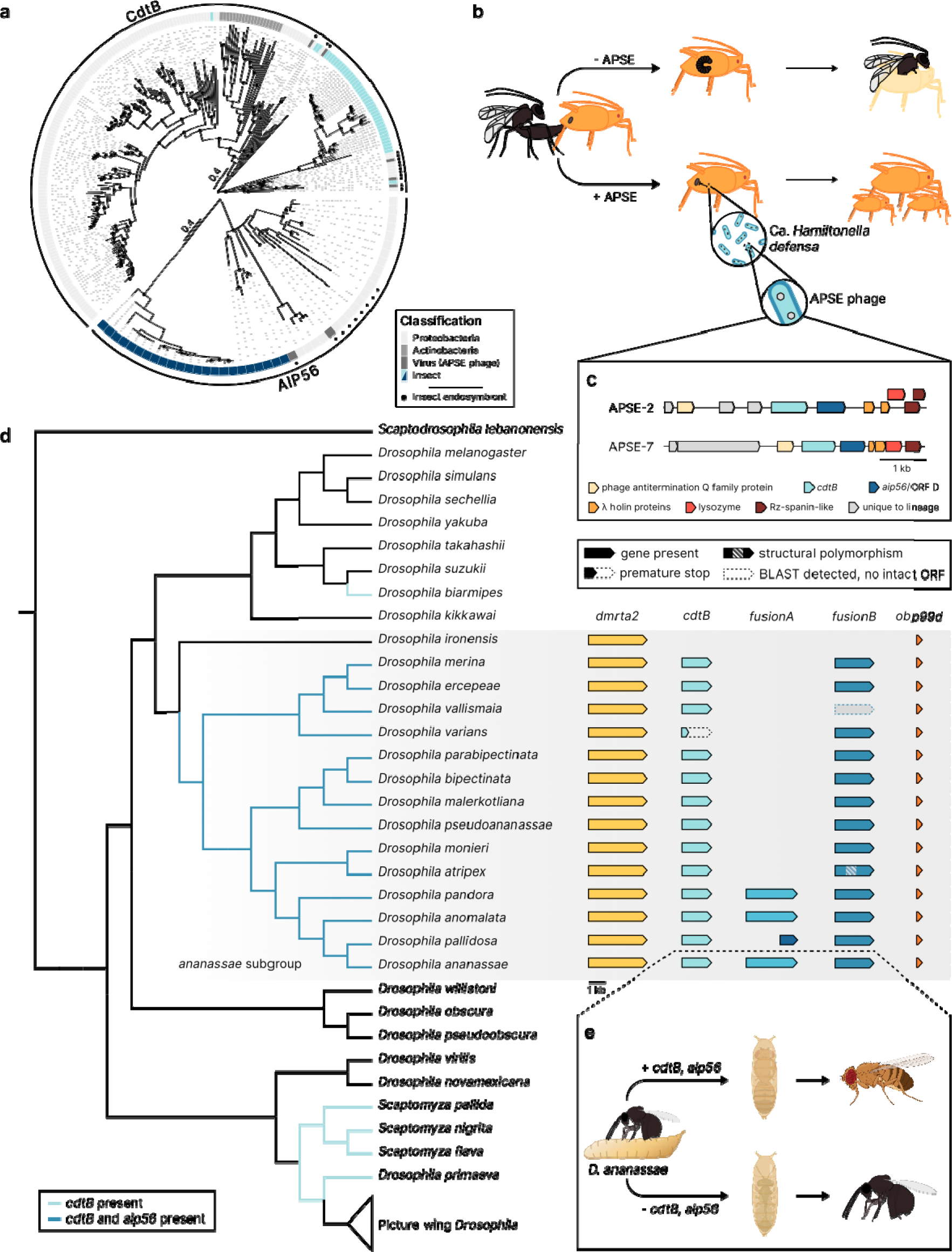
*cdtB* and *aip56* encode defensive toxins associated with insects and thei endosymbionts. (a) Maximum likelihood protein phylogenies for CdtB and AIP56 proteins. Both trees are midpoint rooted. Nodes with ≥90% bootstrap support are labeled. Highlighted clades indicate *ananassae* subgroup homologs. Scale bar = substitutions/site. (b) Cartoon schematic of the defensive symbiosis between aphids and their secondary endosymbiont, Ca. *H. defensa*, when infected with APSE phage. In the absence of APSE phage, parasitoid wasps readily complete development and hatch out of aphid “mummies.” When aphids carry APSE-infected *H. defensa* strains, they survive and reproduce, transmitting the endosymbionts in their offspring. Figure modified from [42]. (c) APSE phage toxin cassette regions encode diverse toxins, including syntenic copies of *cdtB* and *aip56* (ORF D). Overlapping ORFs are staggered for clarity. (d) Distribution of *cdtB* and *aip56* in nuclear genomes of the Drosophilidae (left) and synteny analysis of *cdtB* and *cdtB::aip56 fusion* proteins in the *ananassae* subgroup (right). Species tree topology based on previously published topologies^36,79,82^. Branch lengths are arbitrary. Colored branches are inferred from previously reported maximum likelihood calculations^36^ and maximum parsimony based on synteny analysis. (e) Cartoon schematic of horizontally transferred *cdtB* and *cdtB::aip56* fusion genes in parasitoid defense in *D. ananassae.* Panel (e) was partially created with BioRender.com.

While *cdtB* and *aip56* play a key role in parasitoid wasp defense in diverse insect systems, little is known about how these genes are evolutionarily co-opted into immune roles without intoxicating the host. Here, we experimentally retraced^45^ the HGT event that introduced *cdtB* and *aip56* into a common ancestor of the *ananassae* species subgroup by generating transgenic *D. melanogaster* lines that encode these genes. Using gain-of-function studies in this naïve background, we found that *fusionB* was sufficient to promote fly survival and parasitoid wasp mortality when expressed in immune tissues. We further demonstrated that FusionB is a novel secreted nuclease that functions in humoral immunity. FusionB also induced growth defects when expressed constitutively in flies and in yeast, underscoring its cytotoxicity across diverse taxa. Our study shows how phage-encoded defense systems can contribute to the rapid evolution of animal innate immune defenses via HGT, despite the risk of autoimmunity.

## Results

### *cdtB* and *fusion* genes are evolutionarily conserved among *Drosophila ananassae* subgroup species

We previously discovered the horizontal transfer of *cdtB* and *aip56* from an APSE phage ancestor into the drosophilid lineage. While there have been at least three independent transfers of *cdtB* in drosophilids, *aip56* appears restricted to the *ananassae* subgroup (Figure 1D). To elucidate the evolutionary history of these genes within the subgroup, we surveyed 8 new *ananassae* subgroup genomes for the presence of *cdtB* and the *fusion* genes. We found *cdtB* and at least one copy of the *fusion* gene in all sequenced *ananassae* group species except for *D. ironensis*, which lacked either gene, and *D. vallismaia*, which appeared to have lost its *fusion* gene (Figure 1D, Figure S1). *cdtB* and *fusion* genes were syntenic in all *ananassae* subgroup species (Figure 1D), and individual gene trees largely recapitulated the topology of the species phylogeny (Figure S2). The insect homologs of CdtB were sister to those found in insect endosymbionts and phages that infect them, including APSE-2 and APSE-7. *ananassae* subgroup homologs of AIP56 formed a long branch stemming out from a clade that contained homologs from insect endosymbionts and other insectine homologs of AIP56 (Figure S2). These data suggest a single horizontal transfer event of *cdtB* and *aip56* from an APSE phage ancestor to the common ancestor of all *ananassae* group species following their divergence from the *D. ironensis* lineage.

CdtB orthologs in all *ananassae* subgroup species retained all functionally important catalytic residues^39^, except in the case of *D. varians*, which had a premature stop codon early in its second exon (Figure S3). Splice junctions were conserved in all *ananassae* subgroup orthologs except for *D. parabipectinata*, which had an insertion at the end of its second exon (Figure S3).

Annotation of new genome sequences also suggested that the *fusion* gene was duplicated in the last common ancestor of the *ananassae* species complex. Species in this complex each encoded two copies of the *fusion* gene, except in *D. pallidosa*, which encoded an *aip56* homolog at the locus where *fusionA* was encoded in sister species (Figure 1D). The Fusion orthologs in species that only encode one copy of the gene were synapomorphic with FusionB (Figure S4). Splice junctions were largely conserved across *fusionA* and *fusionB* paralogs and orthologs (Figure S4). AlphaFold predictions of the Fusion toxins had a three domain structure, similar to other short-trip AB toxins (Figure S5A)^46^. The domains with homology to CdtB and AIP56/ORF D were connected by a 171-amino acid “middle region,” which had no homology to any known protein. This three-domain structure also mirrored that of canonical bacterial cytolethal distending toxins, which form a tripartite complex consisting of CdtB and two other proteins, CdtA and CdtC, that sequester the active site of the CdtB and facilitate its trafficking into the host cell (Figure S5A)^47^. All functionally important residues were conserved in the CdtB domains of the Fusion proteins, with the exception of the Fusion protein encoded by *D. ercepeae* and a short *fusion* allele encoded by *D. atripex*, which is the first polymorphism discovered at this locus in any species (Figure S4, Figure S6). Similarly, there were several conserved cysteine residues directly upstream of the region homologous to APSE AIP56, which may be capable of forming disulfide bonds in the holotoxin^48^ (Figure S4, Fig. S5B). However, other residues described as important to bacterial AIP56 function, such as the pH-sensitive hairpin region or T1-like motif required for membrane association and trafficking^48^, were not detected in *ananassae* Fusion proteins, suggesting alternative mechanisms for cellular intoxication have evolved in this gene family.

### *fusionB* is sufficient for parasitoid defense in *D. melanogaster*

CRISPR null mutant experiments in *D. ananassae* demonstrated that each of the *cdtB* and *fusion* genes contributes to the species’s strong wasp resistance phenotype^38^. However, whether these genes play a direct role in parasitoid response is unclear. To test whether these genes were sufficient to drive the observed anti-parasitoid effect, we generated *D. melanogaster* UAS lines encoding *D. ananassae cdtB*, *fusionA*, and *fusionB,* which recapitulated this HGT event in a naive drosophilid background (Figure 2A). We then used age-matched *GAL4 >* UAS F1 larvae in parasitoid infection experiments.

**Figure 2.**
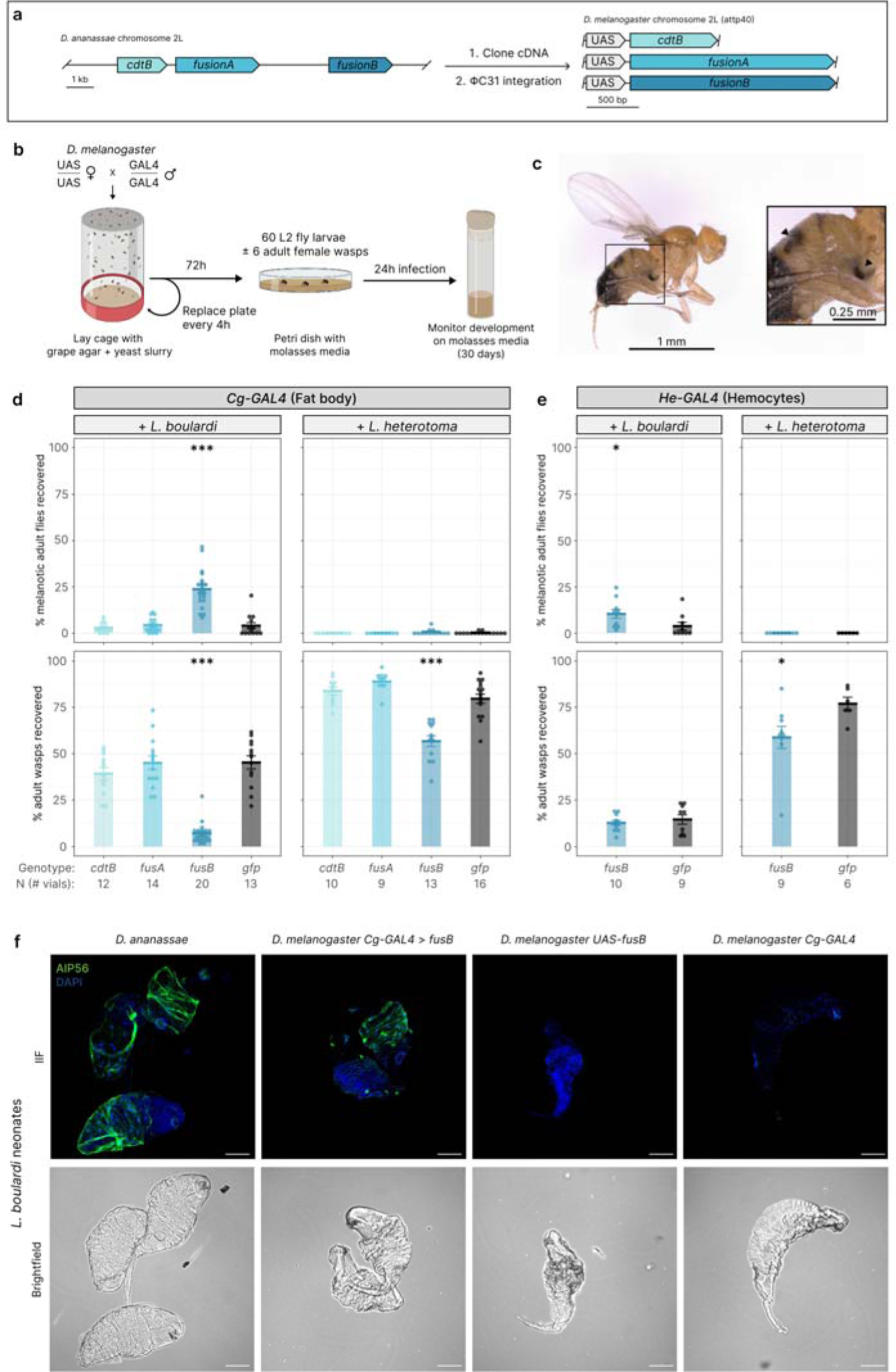
*fusionB* promoted fly survival and inhibited wasp development in *D. melanogaster*. (a) Workflow for generating transgenic *D. melanogaster*. (b) Experimental design for infection experiments. (c) An *L. boulardi (Lb)* infected adult male fly with melanotic cysts, indicating successful neutralization of *Lb* neonates. Inset arrows indicate cysts. (d-e) Parasitization experiment results using fat body-(*Cg-GAL4*, BDSC 7011) and hemocyte-specific (*He-GAL4*, BDSC 8699) GAL4 drivers. Individual data points correspond to the proportion of total larvae infected in an individual vial with each outcome after 30 days (60-67 larvae per vial). Crossbars represent mean ± standard error. (d) *Cg-GAL4* > *fusionB* larvae had higher rates of completing development against specialist wasp *Lb,* but not the generalist *L. heterotoma* (*Lh*). Fewer wasps of either species emerge from *Cg-GAL4* > *fusionB* larvae. *** = p < 0.001, pairwise Wilcoxon rank sum test with Bonferroni correction. (e) *He-GAL4* > *fusionB* larvae experienced a mild increase in adult survivorship against *Lb,* but did not decrease wasp emergence. *Lh* emergence was impaired in *He-GAL4* > *fusionB* flies. * = p < 0.05, Wilcoxon rank test. (f) FusionB protein localized to the wasp neonate serosa in *D. ananassae* and *Cg-GAL4* > *fusionB* larvae, but not in either *D. melanogaster* parent. FusionB was detected via the AIP56 domain (green). Scale bar = 50µm. Panel (b) was partially created with BioRender.com.

Groups of ∼60 second instar larvae were infected either with the *melanogaster* subgroup specialist wasp *Leptopilina boulardi* Lb17, the drosophilid generalist wasp *L. heterotoma* Lh14, or were left as uninfected controls (Figure 2B). The wasp species differ in their infection strategies. *L. boulardi* is the less virulent of the two, largely employing a “passive” mechanism to evade host immune responses by attaching their eggs to host tissues^16^. *L. heterotoma* deploys potent venoms that destroy existing hemocytes and inhibit further hemocyte proliferation to suppress host immune responses^16^. Following a 24 hour infection period, wasps were removed and fly larvae were moved to vials, which were screened for the emergence of adult flies and wasps (Figure 2B). To confirm that surviving adult flies successfully surmounted wasp challenge, we screened adult flies that emerged from wasp-infected vials for melanotic cysts, which form following the native encapsulation reaction that *D. melanogaster* uses in defense against these wasps^24^ (Figure 2C).

Strikingly, *fusionB* showed strong anti-parasitoid effects when expressed from the *Cg-GAL4* driver, which closely replicated the native fat body and hemocyte expression patterns observed in *D. ananassae ^49^*. Against the specialist wasp *L. boulardi, Cg-GAL4 > fusionB* larvae had a higher proportion of adult fly emergence (23.3 ± 10.0%, mean ± standard deviation) vs. the *gfp* control (3.7 ± 5.1%, p = 0.00006, pairwise Wilcoxon rank sum test with Bonferroni correction), and wasp emergence was reduced ∼7-fold (p = 0.000015, pairwise Wilcoxon rank sum test with Bonferroni correction) (Figure 2D, Tables S1-2). Adult flies rarely successfully developed in any genetic background when challenged with *L. heterotoma*, but there was a substantial decrease in wasp emergence from *Cg-GAL4 > fusionB* flies (57.0 ± 10.4%) relative to the *gfp* control (79.8 ± 9.9%, p = 0.00027, pairwise Wilcoxon rank sum test with Bonferroni correction) (Figure 2D, Tables S1-2). We observed no detectable effects of *cdtB-* or *fusionA-*expressing flies versus the *gfp* controls on fly or wasp emergence when challenged with either wasp species (Figure 2D, Tables S1-2). Thus, *fusionB*, but not *cdtB* or *fusionA*, is sufficient to drive the anti-parasitoid phenotype.

The native parasitoid response in *D. melanogaster* is driven by differentiation and subsequent encapsulation of developing parasitoids by specialized hemocytes called lamellocytes^50^. Since F1 larvae in our system retained their cellular anti-parasitoid response, we determined the relative role of fat body *fusionB* vs. hemocyte *fusionB* expression in mediating the anti-parasitoid response we observed using the *Cg-GAL4* driver. Accordingly, we crossed UAS flies to the *He-GAL4* driver (BDSC 8699), which expresses specifically in all hemocytes^51^. Lamellocytes still formed in *He-GAL4 > fusB* larvae following wasp infection (Figure S7), and hemocyte-specific expression did not affect adult survivorship compared to *gfp* controls (Figure S8).

We observed a weaker wasp resistance phenotype using the *He-GAL4* driver than with the *Cg-GAL4* driver (Figure 2E). There was a small increase in adult fly emergence in *He-GAL4 > fusionB* larvae (10.7 ± 7.5%) vs. *gfp* control larvae (3.7 ± 5.8%, W = 73, p = 0.02349, Wilcoxon rank sum test) when challenged against *L. boulardi*, but there was no decrease in wasp emergence vs. the control (W = 38, p = 0.5954, Wilcoxon rank sum test) (Figure 2E, Tables S1-2). When challenged with *L. heterotoma,* no adult flies emerged, but there was a similar decrease in wasp emergence in *He-GAL4 > fusionB* larvae (59.4 ± 18.4%) vs. *gfp* control larvae (76.7 ± 8.6%, W = 47.5, p = 0.018, Wilcoxon rank sum test) compared to that observed in *Cg-GAL4* > *fusionB* larvae (Figure 2E, Tables S1-2). This result was striking given that *L. heterotoma* toxins lyse host hemocytes, perhaps indicating a role for “free” FusionB toxin in the host hemolymph in mediating the anti-parasitoid effect. Therefore, *fusionB* expression in hemocytes alone did not fully recapitulate the strong anti-parasitoid effect observed in the *Cg-GAL4* crosses, highlighting a crucial role for the fat body in mediating this defense response.

Notably, in all experiments, ≥ 80% of fly larvae reached pupariation (Figure S9, Table S1). In uninfected controls, nearly every larva that pupariated successfully developed into an adult (Figure S8B). In the infected groups, many flies were arrested in the pupal stages, resulting in a “stalemate” where neither an adult fly nor an adult wasp eclosed from puparia (Figure S10). *fusionB* expression resulted in more stalemates on average for either wasp when expressed by the *Cg-GAL4* driver, and more stalemates against *L. heterotoma* for the *He-GAL4* driver (Figure S10A-B, Tables S1-2). There was variation in the timing of pupal developmental arrest in infected flies. Some arrested before any adult structures formed and some arrested as pharate adults (Figure S10C). We conclude that *fusionB* expression allowed wasps to progress somewhat in development, resulting in scenarios in which neither the wasp nor the fly completed development.

To determine whether and to what extent FusionB protein interacts directly with developing parasitoids, we performed indirect immunofluorescence (IIF) assays on *L. boulardi* neonate larvae isolated from *Cg-GAL4* > *fusionB* larvae. The AIP56 domain of FusionB was detected in wasp serosal cells in a similar fashion to the “toxic tunic” we previously observed in *D. ananassae* (Figure 2F), indicating that our gain-of-function UAS-GAL4 system recapitulated the native system. We did not detect any FusionB protein on wasp neonates isolated from either the *UAS*-*fusB* or *Cg-GAL4* parents, indicating the reaction was specific to FusionB and not to native *D. melanogaster* or *L. boulardi* proteins (Figure 2F). These results demonstrate that *fusionB* directly antagonized native parasitoid wasp neonates of *D. melanogaster in vivo* and contributed to enhanced survivorship in flies that express the toxin.

### FusionB is a secreted DNase

The large effect of *fusionB* in promoting fly survivorship against *L. boulardi* and repressing wasp development against either wasp when expressed from the fat body *GAL4* driver suggests that *fusionB* encodes a humoral immune factor. To test this hypothesis, we performed western blots on larval fat bodies and on cell-free hemolymph to monitor whether FusionB was secreted into the hemolymph from the fat body. *Cg-GAL4* > *fusionB* flies phenocopy the native fat body expression of FusionB we observed in *D. ananassae* (Figure 3A, Figure S11). Under reducing conditions, we observed appreciable amounts of both the ∼70kDa full length FusionB holotoxin and a ∼45 kDa AIP56 polypeptide, the result of FusionB processing, in the fat bodies of both species. We also detected FusionB protein in cell-free hemolymph of both *D. ananassae* and *Cg-GAL4* > *fusionB D. melanogaster* larvae (Figure 3B), indicating that FusionB protein was secreted from the cells in which it is produced. The major protein species in cell-free hemolymph in either sample was the ∼45kDa fragment, indicating that the majority of FusionB protein is processed once it reaches the host’s hemolymph. This proteolytic nicking is characteristic of AB toxins, including AIP56^46^.

**Figure 3.**
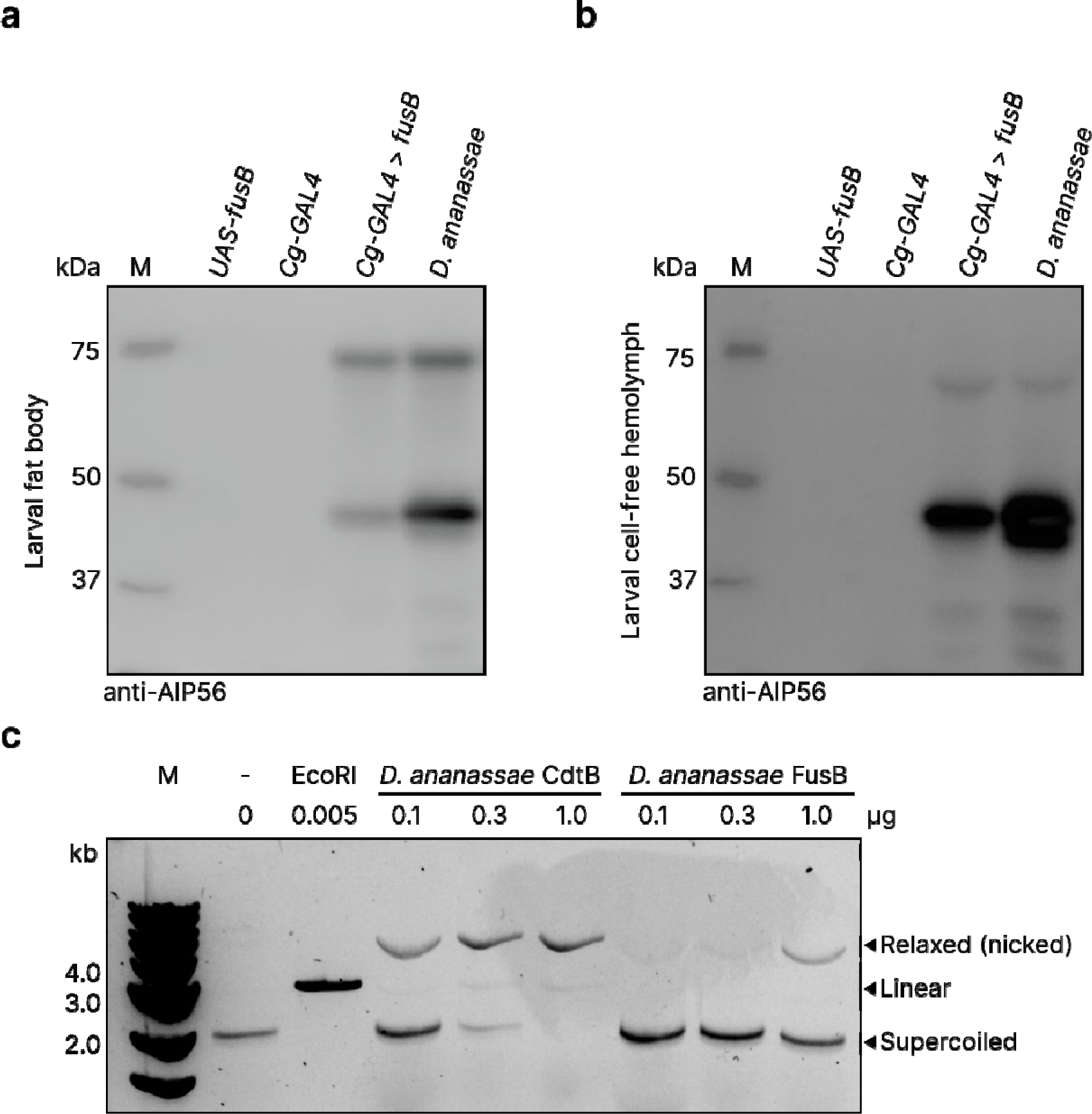
FusionB was secreted from the fat body and functioned as a nickase. (a-b) Western blot analysis of larval fat bodies (a) and larval cell-free hemolymph (b) targeting the AIP56 domain of FusionB under reducing conditions. (a) The full length FusionB holotoxin and its processed AIP56 domain were detected in the fat bodies of *D. ananassae* and *Cg-GAL4 > fusionB D. melanogaster* larvae. 30ng total protein was loaded into each well. (b) The majority of FusionB detected in cell-free hemolymph was in its processed form in both *D. ananassae* and *Cg-GAL4 > fusionB* flies. 15ng total protein was loaded into each well. (c) Plasmid relaxation assay indicated that FusionB had nickase activity *in vitro*.

Based on structural predictions and conservation of key catalytic residues, we hypothesized that FusionB CdtB retains its nuclease activity and tested whether FusionB has such activity *in vitro.* We found that FusionB had nickase activity *in vitro,* similar to *D. ananassae* CdtB (Figure 3C)^36^. Taken together, these results demonstrate that FusionB is secreted from the fat body into host hemolymph where it can act humorally, and the enzyme retained its toxic nuclease activity *in vitro*.

### Constitutive *fusionB* expression is lethal

*cdtB* and *aip56* toxins originated in bacteria and cause serious damage to eukaryotic cells^39,40^. We hypothesized that these toxins must be carefully expressed in recipient *Drosophila* lineages to avoid autoimmunity. To address this working hypothesis, we crossed balanced UAS *D. melanogaster* flies to balanced *Actin-GAL4 D. melanogaster* flies, which resulted in constitutive expression of the transgene across all life stages and tissues in F1 progeny. We hypothesized that if constitutive expression of these transgenes were toxic to the fly, we would observe fewer F1 flies without the balancer phenotype than expected via Mendelian genetics. Strikingly, these *D. melanogaster* flies survived constitutive expression of *D. ananassae cdtB* and *fusionA* (Figure 4A, Table S3). However, nearly 100% of flies from the *Actin-GAL4* x *fusionB* crosses had the balancer phenotype, indicating constitutive *fusionB* expression caused complete developmental failure (Figure 4A, Table S3). We confirmed CdtB and FusionA protein were expressed in adult F1 flies via western blot, validating these results were not due to failure of transgenic flies to produce the protein (Figure S12). Thus, *fusionB* is unique among insect-associated *cdtB* homologs in its toxicity to *D. melanogaster*.

**Figure 4.**
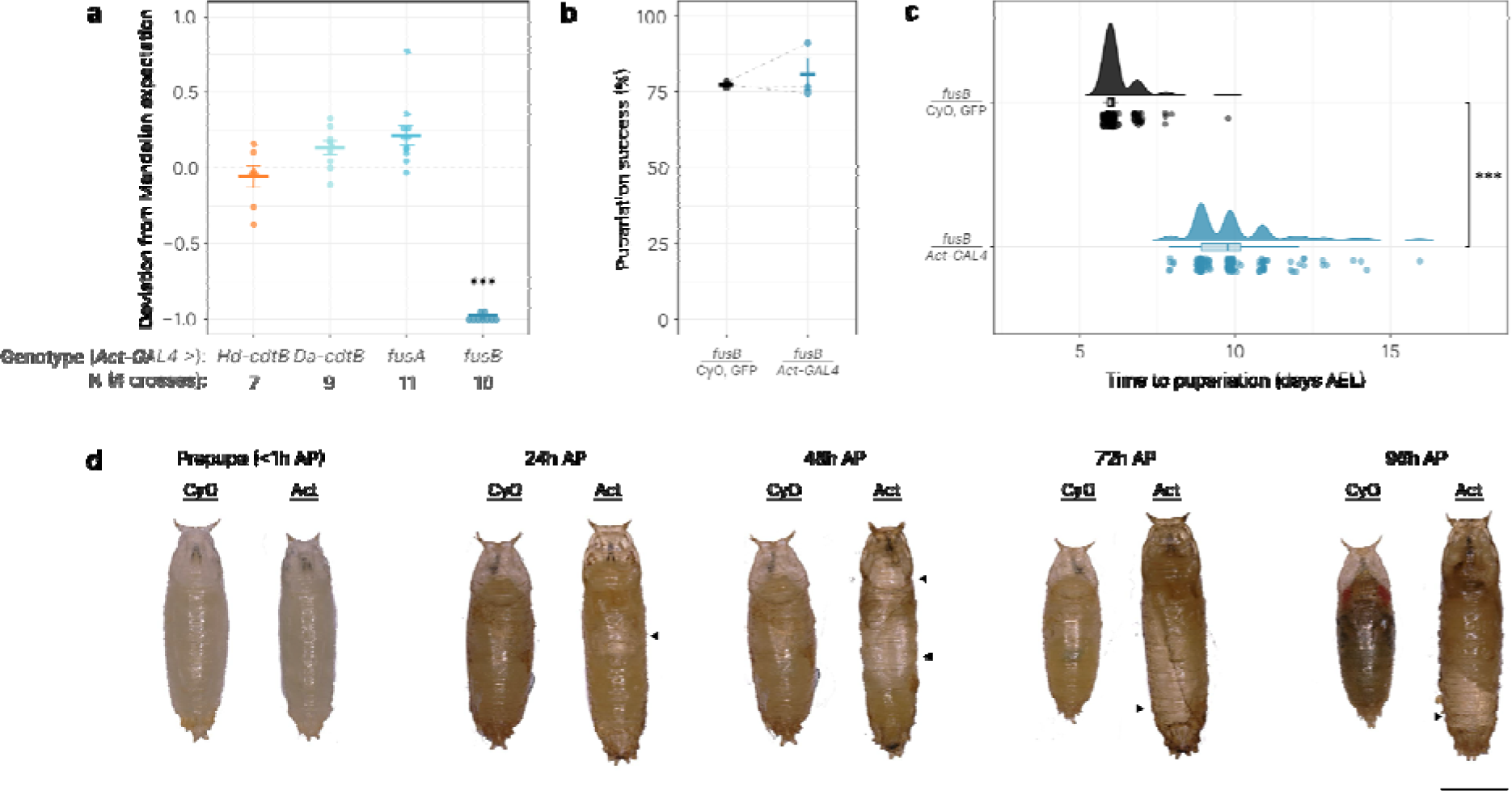
*fusionB* retained its toxicity in *D. melanogaster*. (a) Constitutive expression of *fusionB*, but not *Hd-cdtB, Da-cdtB*, or *fusionA,* from the *Actin-GAL4* driver (BDSC 4414) arrested development in *D. melanogaster*. *** = p < 0.001, one-sided exact binomial test (H_a_: observed < expected). Crossbars indicate mean ± standard error. (b) Pupariation success of *fusionB/Act-GAL4* and *fusionB*/CyO, GFP siblings did not differ (N = 3 crosses, p = 0.5644, X-squared = 0.33213, df = 1). Crossbars indicate mean ± standard error. Grey dashed lines compare pupariation rates of siblings from the same cross. (c) Constitutive expression of *fusionB* disrupted timing of pupariation (*fusionB/Act-GAL4* n = 149 larvae, *fusionB/*CyO, GFP n = 146 larvae). (*** = p < 0.001, W = 21676, two-sided Wilcoxon rank sum test). For boxplot: center line, median; box margins, first and third quartile; whiskers, 95% confidence interval. Outliers are not shown on boxplots. AEL = after egg lay. (d) Age-matched pupae of *fusionB*/CyO, GFP (CyO) and *fusionB/Act-GAL4* (Act). Constitutive expression of *fusionB* caused developmental failure at the ‘gas bubble’ stage of pupariation. Arrows indicate aberrant gas bubble presence in *fusionB/Act-GAL4* pupae. AP = after pupariation.

We also determined if *cdtB* and *fusionA* had become non-toxic through evolution following their incorporation into insect nuclear genomes by generating a UAS-*H. defensa cdtB D. melanogaster* line. The expected Mendelian proportions of balancer and non-balancer siblings were recovered from *Actin-GAL4 x* UAS-*H. defensa cdtB* crosses, indicating that *H. defensa cdtB* was broadly non-toxic to *D. melanogaster* as well (Figure 4A). We additionally attempted to generate UAS-*E. coli cdtB D. melanogaster*, but failed to recover transformants after several attempts at different landing sites, DNA concentrations, and growth temperatures, suggesting that even small titers of *E. coli* CdtB are toxic to insects. These data suggest that *H. defensa cdtB* has coevolved with insects, while *E. coli cdtB* has not.

Having established that *Actin-GAL4* > *fusionB* flies failed to complete development, we next determined at what stage the developmental failure occurred. We crossed homozygous *UAS*-*fusionB* flies to *Actin-GAL4*/*CyO, gfp* flies, which allowed us to separate balancer siblings from experimental siblings in early larval stages using a GFP signal. We then monitored the development of age-matched siblings. The siblings largely match each other developmentally until the late third instar stage. While siblings eventually pupariate with similar levels of success (p = 0.5644, X-squared = 0.33213, df = 1) (Figure 4B), pupariation time was severely delayed in *Actin-GAL4* > *fusB* siblings (9.86 ± 1.30 days vs. 6.2 ± 0.53 days for balancer siblings, W = 77.5, p < 2.2e-16, Wilcoxon rank sum test) (Figure 4C). The distribution of time to pupariation was much wider for *Actin-GAL4 > fusionB* larvae compared to balancer siblings (Figure 4C), and wandering third instar larvae were observed in vials as long as 19 days after egg lay. Within 24 hours of pupariation, *Actin-GAL4 > fusionB* flies experienced catastrophic developmental arrest during metamorphosis, likely at the gas bubble stage of pupariation, after which they failed to form any adult structures (Figure 4D, Figure S13). Therefore, constitutive expression of *fusionB* caused severe growth defects typically resulting in mortality, likely in a cell-type- or life-stage-specific manner.

### Growth defect phenotype found in flies is mirrored in yeast

Our experiments focused on the function of *cdtB* and the *fusion* genes in *D. melanogaster*, an organismal context similar to the one in which they evolved. We next determined whether *cdtB* or *fusionA* were generally benign in other eukaryotic backgrounds, or whether there were genetic factors in the drosophilid lineage that abrogate toxicity. To do so, we cloned several bacterial and insect *cdtB* and *aip56* homologs, as well as *D. ananassae fusionA* and *fusionB,* into galactose inducible expression vectors for *Saccharomyces cerevisiae*, a eukaryote that is evolutionarily highly divergent from fruit flies and is unlikely to have encountered these toxins natively.

We performed a dilution series to determine the toxicity of each gene in yeast and compared growth of each strain on galactose-containing media (induced condition) and glucose-containing media (non-induced control). All strains grew comparably on glucose-containing media, indicating any growth defect phenotype on galactose was the result of toxin expression (Figure 5A, Figure S14). As previously reported, *E. coli cdtB* induced severe growth defects in *S. cerevisiae*^52^*. H. defensa cdtB* also strongly inhibited growth, as did APSE-7 *aip56*, though to a lesser extent than those of the bacterial CdtB homologs (Figure 5A). Strikingly, yeast expressing *D. ananassae cdtB* grew equivalently to the empty vector control, suggesting broad non-toxicity of *D. ananassae* CdtB to eukaryotes (Figure 5A). This non-toxicity holds true even when expressing *cdtB* homologs from drosophilid lineages that possess only one copy of *cdtB* (Figure S15), suggesting this is not a feature of subfunctionalization after duplication events. Induction of each *fusion* gene induced a growth defect in yeast, unveiling cryptic toxicity for *fusionA* (Figure 5A). We conclude that the FusionA and FusionB toxins retained ancestral toxicity, while single-copy insectine CdtB evolved to become benign in a eukaryotic context.

**Figure 5.**
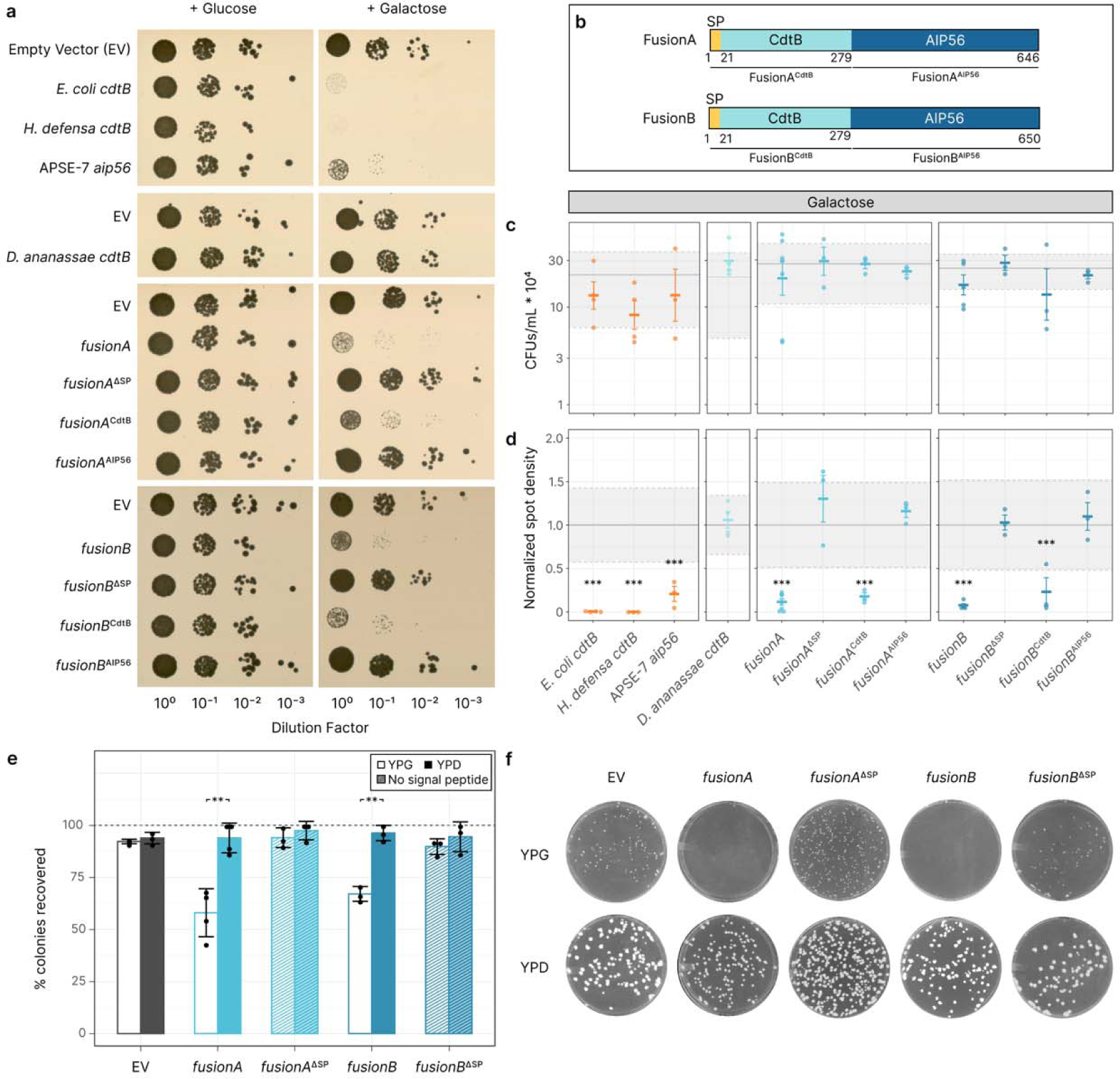
*D. ananassae* Fusion proteins caused growth defects in yeast. (a) Representative images of spot assays on CSM-Leu media containing either 2% glucose (non-inducing, left) or 2% galactose (inducing, right). (b) Cartoons of protein domains for each Fusion protein. (c) Colony forming unit (CFU) counts for each strain on galactose media. Crossbars indicate mean ± standard error. Grey boundaries indicate the mean (solid line) and 95% confidence interval (grey rectangle) of CFU counts for empty vector controls used in experiments with each strain. (d) Spot densities for each strain on galactose media, normalized to the average spot density of empty vector controls used in experiments with each strain. Crossbars indicate mean ± standard error. The grey boundaries indicate the mean (solid line) and 95% confidence interval (grey rectangle) of spot densities for empty vector controls. *** = p-adj < 0.001, Tukey’s test for pairwise comparisons. (e) Percentage of colonies recovered after 48 hours from replica plating strains onto YPG (open bars) and YPD (closed bars). Error bars represent standard deviation. * = p-adj < 0.05, ** = p-adj < 0.01, Student’s t-test with Bonferroni correction. (f) Representative images of YPG and YPD replica plates for each strain at 48h post-replication.

We next created a series of truncation mutants to determine which domains were essential to Fusion protein toxicity in yeast (Figure 5B). Removal of the signal peptide from each Fusion toxin completely ablated its toxicity, suggesting the toxin must be secreted in order to exert its effects (Figure 5A). The CdtB domain (residues 1-279) of each Fusion protein also induced a growth defect, but the AIP56 domain (FusionA residues 280-646 and FusionB residues 280-650) of each Fusion protein grew comparably to the empty vector control (Figure 5A). Therefore, the CdtB domain of the Fusion proteins underlies its toxicity in yeast, not the AIP56 domain.

Curiously, none of the toxins completely inhibited growth upon expression, as the number of colony forming units (CFUs) did not differ from the empty vector control for any strain (Figure 5C). Rather, colonies were much smaller than those of the empty vector controls in strains that expressed toxic products (Figure 5D), indicating the toxins slow growth rather than killing cells outright.

The small colony phenotype we observed in strains expressing the full length Fusion toxins is reminiscent of *S. cerevisiae petite* mutants, which are deficient in their capability to undergo aerobic respiration^53^. This phenotype suggests that the Fusion toxins may impair mitochondrial function in yeast. We tested mitochondrial function by replica plating yeast grown on galactose plates onto YPG media, in which glycerol requires yeast to undergo aerobic respiration, and YPD media, in which dextrose can support yeast growth via fermentation alone. If mitochondrial function was inhibited in yeast expressing the Fusion toxins, we expected to observe growth defects on YPG, but growth should be supported on YPD. After 48 hours, approximately 30% fewer colonies were recovered on YPG for yeast that expressed the full length Fusion toxins compared to the empty vector control (Figure 5E). The colonies that grew on YPG were markedly smaller than those of the empty vector control (Figure 5F). This phenotype was rescued in strains that expressed the truncated Fusion proteins (Figure 5E-F). Growth on YPD was comparable to that observed in the empty vector and truncated Fusion proteins (Figure 5E-F). These results are consistent with the Fusion toxins acting on the mitochondria.

## Discussion

Chronic, long-term coevolution with macroparasitic animals has played a preeminent role in shaping genome-wide patterns of local adaptation in humans and may even contribute to widespread autoimmunity^12^. Unraveling the mechanisms driving these patterns is challenging however, particularly using *in vivo* studies involving macroparasitic animals of humans. The *Drosophila* lineage and the parasitoids that attack it provide an excellent model for such studies given the analogous Red Queen-like co-evolutionary dynamics that have unfolded between fly and parasitoid and the availability of the genetic tools of *D. melanogaster*. Here, we used gain of function experiments in *D. melanogaster* to retrace^45^ the horizontal transfer of novel anti-parasitoid effectors that occurred ca. 21 million years ago in the *D. ananassae* lineage. Immune tissue expression of one of these toxins, *fusionB*, was sufficient to promote fly survival and parasitoid wasp death. FusionB was secreted into host hemolymph where it interacted with developing wasp neonates. Constitutive expression of *fusionB* delayed pupariation and eventual developmental arrest at early pupal stages, indicating latent costs of autoimmunity. Finally, expression experiments in yeast demonstrated that the Fusion toxins evolved eukaryotic signal peptides essential to their function and exerted toxic effects via the CdtB and not the AIP56 domain.

To our knowledge, FusionB is the first toxic humoral anti-parasitoid factor described in *Drosophila*. Previously characterized anti-parasitoid immune strategies in *Drosophila* are largely cell-mediated, where wasp infection causes a hematopoietic burst followed by differentiation of specialized hemocytes that subsequently encapsulate wasp eggs and larvae^24,27,44^. However, many species of *Drosophila* fail to encapsulate embryos^25^, and therefore must rely on other mechanisms to neutralize parasitoids. While there is extensive literature describing humoral responses to parasitoid attack, these generally refer to compounds that do not have clear anti-parasitoid roles, such as the production of anti-microbial peptides (AMPs)^16^ that are instead likely to be activated in response to the introduction of microbes during oviposition by wasps or reactive oxygen species generated by prophenoloxidase enzymes produced by specialized hemocytes. Other fat body factors implicated in parasitoid defense in *D. melanogaster* influence hemocyte production (e.g., Edin)^54^, or encourage recruitment of hemocytes to the wasp egg (e.g., Lectin24A)^55,56^. Similar to aphids, some *Drosophila* species have adopted facultative symbioses with bacteria like *Spiroplasma,* which express RIPs to protect against parasitoid challenge^21,57^. However, these symbioses often come with severe fitness costs either due to the inability of the bacterial toxins to discriminate between host and parasitoid targets^58^ or through other toxic activities of the endosymbiont, such as male-killing^57^. Thus, our study illuminates the recruitment of *cdtB* and *aip56* into the native humoral immune system of some *Drosophila* as a potent mechanism by which they defend against parasitoid wasp challenge.

Notably, insect CdtB and the Fusion toxins do not appear to kill cells outright. CdtB itself had no clear toxic effect in yeast, flies, or against wasps, despite retaining potent nuclease activity (Figure 3C). This points to its ancestral domestication to a eukaryotic environment. *cdtB* knockouts adversely affected *D. ananassae* resistance to a diverse panel of parasitoid wasps^38^, suggesting that while CdtB does not interact directly with parasitoid wasps, it may play an indirect role in facilitating wasp resistance through unknown mechanisms. In contrast, the toxicity of the Fusion paralogs varied based on the cellular and regulatory context. FusionA was broadly non-toxic to flies and wasps but exerted a potent growth defect in yeast. FusionB was broadly toxic to both wasp species tested, caused severe growth defects in *D. melanogaster* when expressed constitutively, and arrested growth in yeast. We hypothesize that the Fusion toxins act as weak effectors that primarily slow down cellular growth and proliferation, which opens a window for other host immune effectors to act against these enemies (Fig. S16). Such a model has been proposed for bacterial cytolethal distending toxins^59^, and could be extended to host-parasitoid systems, in which outcomes are strongly influenced by the host’s developmental stage^60,61^.

Much remains to be understood about the specific mechanism by which FusionB exerts its toxic effects. We hypothesize that the Fusion proteins interact with host factors or receptors that confer cell-type specific toxicity and that the toxin exerts its effects on mitochondrial, rather than nuclear, DNA. Cell-type specificity is a recurrent theme among bacterial AB toxins, perhaps best exemplified by botulinum toxin^62^. Our constitutive expression fly model suggests that FusionB requires interaction with particular host factors to become toxic. This is because broad, non-specific toxicity would likely arrest development in embryonic stages^63^, and *fusionB* expression in immunogenic tissues did not result in any sort of growth defect. Similarly, Fusion toxin localizes specifically to *Leptopilina* serosal tissue, which is shed at the first instar stage^60^ and differentiates into “trophic cells”^26^ that are likely teratocytes heretofore unstudied in *Leptopilina* species^64^. Though little is known about these trophic cells or teratocytes, they often play key roles in both extracting nutrients from the host, releasing AMPs, and manipulating host development^64^. Elucidating cell-type specificity can illuminate the mechanisms by which these genes serve as potent toxins against their targets without causing host auto-immunity.

Furthermore, the growth defect phenotype upon expression of *fusionB* in flies and yeast suggest that mitochondrial DNA, rather than nuclear DNA, is damaged. Yeast expressing the *fusion* toxins exhibited a strong growth defect on glycerol, a non-fermentable substrate, indicative of mitochondrial dysfunction. Similarly, our constitutive expression fly model phenocopies mtDNA maintenance mutants, which experience extended larval stages^65^ and typically die at late larval or early pupal stages^65,66^. As such, identifying the molecular mechanisms by which FusionB intoxicates cells and its subcellular targets stand as promising directions for future research.

Our work demonstrates that HGT has been a powerful mechanism by which animals can rapidly expand their innate immune repertoires, even against macroparasitic enemies like parasitoid wasps. Finally, this study shows that phage-derived components that have played such a key role in shaping prokaryotic immune systems also underlie key innovations in animal immunity.

## STAR Methods

### Protein phylogenies

CdtB homologs were found via a BLASTP search^67^ against the NCBI clustered nr database using APSE-2 CdtB (YP_002308521.1) as a query. Full length CdtB sequences were downloaded and run through cd-hit v 4.8.1^68^ with a similarity threshold of 90% to pare down the number of sequences represented in the phylogeny. *D. ananassae* group CdtB sequences were then concatenated to the BLAST sequences and aligned using MUSCLE v 5.1.linux64^69^. The alignment was manually inspected in AliView ^70^. The variable N-terminal signal peptides were trimmed, then redundant or short (<50 aa) sequences were removed from the alignment. The remaining sequences were trimmed on the Clipkit webserver using the kpic-gappy algorithm with a 0.9 gap cutoff^71^. The trimmed sequences were then realigned using MUSCLE. AIP56 homologs were found via a BLASTP search using APSE-2 AIP56/ORF D (YP_002308522.1) as a query. The aligned sequences were downloaded, then run through cd-hit with a similarity threshold of 97%. *D. ananassae* group AIP56 sequences were concatenated to the BLAST sequences and aligned using MUSCLE. The alignment was manually inspected in AliView, where redundant sequences were removed and N-terminal variable sequences were trimmed. The remaining sequences were trimmed on the Clipkit webserver using the kpic-smartgap algorithm, then realigned using MUSCLE. Phylogenetic trees were generated on the IQTree Webserver^72^ using the best-fit model as determined by the ModelFinder feature (CdtB: VT+F+G4; AIP56: WAG+F+I+G4). The trees were then visualized using ggtree v 3.10.1^73^ and ggtreeExtra v 1.12.0^74^. Trees were midpoint-rooted using phytools v 2.1.1^75^ and nodes with <50% bootstrap support were collapsed using the di2multi function in ape v 5.7.1^76^.

### Whole-genome alignment and comparative annotation

Previously assembled *ananassae* subgroup genomes^77–79^ were downloaded from NCBI (Table S4). These species represent 16 out of 22 described *ananassae* subgroup species. *D. setifemur* was set as the outgroup to all other taxa listed here. A reference-free whole-genome alignment was constructed with the Progressive Cactus v.2.6.0 software^80^. The phylogenetic relationships inferred by Kim et al. 2023 were supplied to the alignment software as the guide tree for the alignment. The Comparative Annotation Toolkit software^81^ v2.2.1, was used to annotate all other taxa in the alignment using the protein-coding gene *D. ananassae* NCBI RefSeq annotations as a reference. The *cdtB* and *fusion* loci were then manually inspected for accuracy. *D. ananassae cdtB* and *fusion* gene boundaries were used as templates to correct mis- or unannotated loci.

### Species tree construction

The species tree topology was based on previously published topologies. For the *ananassae* group, topologies were based primarily on Matsuda et al. (2009)^82^. New taxa (i.e. *D. anomolata*) were placed based on topologies predicted from whole genome alignments^79^. Branches are colored based ancestral state predictions elucidated previously^36^.

### Amino acid alignments

Predicted amino acid sequences for CdtB and Fusion proteins were extracted using the Translate function on Geneious Prime 2023.1.2. Protein sequences were then aligned using Mafft v 7.490^83^ as implemented in Geneious. Residue conservation was scored using the BLOSUM62 matrix.

### Splice site prediction for *D. ananassae cdtB* and *fusion* genes

Annotation predictions generated using the Comparative Annotation Toolkit software were used to predict splice sites in the new *ananassae* group genomes. In cases where the toolkit predicted additional junctions or where junction predictions resulted in a truncated protein, the Berkeley Drosophila Genome Project Splice Site Prediction by Neural Network tool^84^ was used to predict alternate splice junctions. Manually curated gene model predictions retained as much of the *D. ananassae* gene structure as possible.

### AlphaFold structural predictions

The sequences for the *E. coli* strain B574 CDT complex were downloaded from GenBank (CdtA: OSK27480.1; CdtB: OSK27479.1; CdtC: WP_000825549.1). The solved *Haemophilus ducreyi* CDT crystal structure^47^ (PDB 1SR4) was used as a template for Alphafold Multimer^85^ as implemented through ColabFold v 1.5.5^86^ in March 2024. The sequences for *D. ananassae* FusionA (XP_044570248.1) and FusionB (XP_032305712.2) proteins were submitted to the ColabFold v 1.5.5 notebook in February 2024. Structures were generated using the default Alphafold settings^87^. The rank 1 structures for each protein were visualized in ChimeraX v 1.7^88^.

### Disulfide bond site predictions

FusionA and FusionB structures were screened by eye for cysteine residues in close proximity in the 3D structure of the protein. No such pairs were observed in FusionA structural predictions. FusionB had a pair between C301 and C448. To calculate the distance between these residues, the sulfur atoms were selected in ChimeraX v 1.7^88^ and the distance command was used to calculate the distance between them.

### D. atripex genotyping

Single legs from D. atripex strains CS Ng, V251, and Y226 were dissected and incubated in Buffer A (100 mM Tris-Cl (pH 7.5), 100 mM EDTA, 100 mM NaCl, 0.5% SDS) + 5% Proteinase K (NEB) at 50°C overnight to isolate genomic DNA. 4 adult males for each strain were used as samples. The reactions were deactivated by heating them to 95°C for 10 minutes. 2µl of the resulting DNA prep were used as template in a PCR using the following primers. PCR products were run on a 1% TAE agarose gel stained with SYBR Safe (Invitrogen) at 120V for 25 minutes. The gel was visualized on a ChemiDoc MP Visualization System (BioRad). Each lane represents a single individual.

### Insect husbandry

All insects used in the study were reared at 24°C, 60% relative humidity, on a 12:12 light:dark cycle, unless otherwise indicated. *Leptopilina* wasp cultures were reared on *D. melanogaster w1118*. Identities of all insect cultures used in the study can be found in Key Resources table.

### Insert generation for cloning

All inserts for transgenic experiments, except for *E. coli*-optimized *E. coli cdtB*, *S. flava cdtB, and D. ananassae cdtB,* were synthesized by Twist Biosciences. *E. coli-*optimized *E. coli cdtB* was synthesized by Genewiz/Azenta Biosciences. Information about all new gene fragments synthesized for this study can be found in Table S5. *S. flava cdtB* was amplified off of a 2B-T plasmid^17^. Intronless *D. ananassae cdtB* was obtained through cDNA isolation using the Protoscript II kit (NEB) from total RNA isolated from *D. ananassae* embryos using the NEB RNA miniprep kit. Gene blocks used to generate *Drosophila* transgenic strains encoding bacterial CdtB copies were codon-optimized for *Drosophila melanogaster.* Bacterial *cdtB* inserts cloned into both flies and yeast did not include their N-terminal signal peptides. Information about all primers used for cloning can be found in Table S6.

### *Drosophila* strain generation

Inserts encoding *E. coli, H. defensa, and D. ananassae cdtB,* and *D. ananassae fusionA and fusionB* were PCR amplified using Q5 polymerase (NEB) and cloned into pUAST::attB^89^ at the NotI restriction site using the HiFi DNA Assembly Kit (NEB). Constructs were sequence validated via Sanger sequencing or Plasmidsaurus whole plasmid sequencing. Sequence verified plasmids were sent to Genetivision Corporation for phiC31 integration at the attP40 (Ch. 2) or attP2 (Ch. 3) sites.

### Parasitization experiment

Homozygous virgin females were collected from UAS lines in groups of ∼80-120 and crossed to 20-30 homozygous GAL4 males. We selected UAS-*gfp* (BDSC 5431) as a control line to express a non-native transgene that we expected to have no impact on wasp development. Crosses were moved to chambers outfitted with grape agar petri dishes supplemented with live yeast to encourage oviposition. Fresh plates were switched every four hours to synchronize larval development. Larvae developed until the late L2 stage (∼60-70 hours), then transferred in groups of 60 into petri dishes containing 600ul molasses fly media. Each petri dish was fitted with a water-soaked piece of Whatman 3 paper on the lid to retain moisture during the experiment. Petri dishes were randomly assigned to treatment groups (no wasp (uninfected), + *L. boulardi*, + *L. heterotoma*). When larvae were ∼72 hours old, 6 mated female wasps were added to dishes in the infected treatments and were allowed to oviposit for 24 hours. Larvae were then transferred to food vials to complete development. Vials were screened for pupation and adult fly and wasp emergence for 30 days post-infection. Wasp infection in surviving adult flies was confirmed by screening for melanotic cysts that form upon encapsulation of wasp neonates. Vials where > 20% of the larvae developed into adults with no visible melanotic cysts were excluded from further analysis as infection was considered incomplete.

### Indirect immunofluorescence assay

Third instar naïve or *L. boulardi* infected (72 h after parasitization) larvae were dissected in Schneider’s medium (Lonza) supplemented with 5% fetal bovine serum (Gibco) and 1 nM 1-phenyl 2-thiourea (Sigma) (CSM) and fat bodies or parasitoids were isolated. The samples were fixed with 2% paraformaldehyde for 10 min, washed three times in PBS (5 min each), and blocked with 0.1% BSA in PBS supplemented with 0.1% Triton X-100 for 20 min. Hemocytes were isolated in CSM on multispot microscope slides (Hendley-Essex), adhered for 1 h, fixed with acetone for 6 min, air dried, and blocked with 0.1% BSA in PBS for 20 min. Following this, the samples were incubated with either anti-AIP56 2H5^38^ or anti-lamellocyte L1^90^ primary monoclonal antibodies in the form of undiluted hybridoma supernatants containing saturating amounts of the specific immunoglobulins. The fat bodies and the parasitoids were incubated overnight at 4°C and the hemocytes for 1 h at room temperature. Next, samples were washed three times in PBS (each 5 min), incubated with the Alexa Fluor 488 goat anti-mouse IgG secondary antibody (Invitrogen, 1:1,000), containing DAPI (2.5 μg/mL) for 45 min, washed three times in PBS, mounted in Fluoromount G medium, and analyzed with an Olympus FV1000 confocal laser scanning microscope or an epifluorescence microscope (Zeiss Axioscope 2 MOT).

### Protein sample preparation and western blot analysis

Fat bodies were isolated in sterile PBS from naïve third instar *D. melanogaster* larvae. Positive control samples were obtained from *L. boulardi-*infected *D. ananassae* 72 h post-parasitization. The extracts were prepared in sample buffer (250 mM Tris pH = 6.8, 35% glycerol, 0.75 mg/mL Bromophenol blue, 9.2% SDS) using a homogenizer, incubated for 1h on ice, then boiled for 5 min and centrifuged at 18,000xg for 5 min. Hemolymph was harvested by bleeding 60 third instar larvae in 150 μl sterile PBS supplemented with 1 nM 1-phenyl 2-thiourea (Sigma), 1mM PMSF (Sigma) and Complete Protease Inhibitor Cocktail (Roche) according to the manufacturer’s instructions. Isolates were centrifuged at 4°C and 500xg for 5 min to remove the hemocytes. Protein concentrations were determined by the Amido Black assay. To generate reduced conditions, 5% β-Mercaptoethanol (β-ME) was added. 30µg fat body and 15µg hemolymph crude protein extracts were loaded on 10% SDS PAGE. Proteins were blotted to polyvinylidene difluoride (PVDF) membrane (Millipore), which was blocked with 5% nonfat milk in Tris-buffered saline (TBS) (10 mM Tris pH 7.5, 150 mM NaCl). Membranes were incubated with the AIP56 specific 2H5 monoclonal antibody^38^ as hybridoma supernatant used in 1:1 dilution with RPMI Medium1640 (Gibco) containing 5% fetal bovine serum (FBS) for 1h. Membranes were then washed three times (10 min each) with TBS containing 0.1% Tween 20, incubated with Polyclonal Goat Anti-Mouse Immunoglobulins/horseradish peroxidase (HRP) (Dako) (1:10,000 diluted in TBS containing 0.1% Tween 20 and 1% bovine serum albumin (BSA)) for 1 h, and washed three times (10 min each) with TBS containing 0.1% Tween 20 and two times in TBS. Reactions were visualized with Immobilon Western Chemiluminescent HRP Substrate (Merck).

### Protein sample preparation and western blot analysis for *Actin-GAL4* crosses

Adult F1 offspring of UAS/*Actin*-*GAL4* crosses were collected at 5-7 days after eclosion. Curly-winged (CyO) siblings were collected for negative controls. Each sample was prepared with 10 whole females and 200 uL of 1X Laemmli Sample Buffer (Bio-Rad) + 5% β-Mercaptoethanol. The flies were crushed with sterile pestles and centrifuged for 1 min at 15,000 rpm twice. The samples were boiled for 5 min, vortexed, and placed on ice for 5 min before a final centrifugation at 15,000 rpm for 1 min. 10 µL of supernatant was loaded on 10% SDS PAGE gel. Proteins were blotted to polyvinylidene difluoride (PVDF) membrane (Bio-Rad), and transfer was confirmed using Ponceau-S staining (0.1% Ponceau S, 5% glacial acetic acid in distilled water). The membrane was blocked for one hour with 5% nonfat milk in Tris-Buffered Saline containing 0.1% Tween20 (TBST) (10 mM Tris pH 7.5, 150 mM NaCl). Primary antibody staining was carried out overnight at 4°C with CdtB-specific 3G9 monoclonal antibody (UAS-*cdtB* x *Actin-GAL4* crosses) or AIP56-specific 2H5 monoclonal antibody^19^ (UAS-*fusionA* x *Actin-GAL4* crosses) diluted 1:10 in fresh blocking buffer. After washing three times (5 min each) with TBST, the membrane was incubated for an hour with Polyclonal Goat Anti-Mouse Immunoglobulin/Horseradish Peroxidase (HRP) diluted 1:2000 in blocking buffer. The membrane was washed three times for 5 min with TBST and visualized with Western Lightning Plus-ECL (Perkin-Elmer).

### Recombinant protein purification

*D. ananassae* CdtB used in this study was purified as described previously^36^. To generate recombinant cdtB-aip56B-[21-650aa] (hereafter referred to as FusionB), DNA encoding c*dtB-aip56B* (excluding the predicted N-terminal 20-mer-long signal peptide) was PCR amplified using Q5 High-fidelity DNA polymerase (NEB) from *Leptopilina boulardi* G486 infected *D. ananassae* larval cDNA library using gene-specific primers listed in Table S6. The PCR product was inserted into the pDONR221 donor plasmid using the Gateway System (Thermo Fisher Scientific, Waltham, MA, USA) and FusionB was subcloned into the pDEST17 destination vector (Thermo Fisher Scientific). All constructs were verified by DNA sequencing. Recombinant FusionB was expressed in SixPack *E. coli* strain^91^ as follows: bacteria were cultured in 50 ml Terrific Broth auto-induction medium (#AIMTB0201, Formedium) supplemented with 100 µg/mL Carbenicillin for 18h, at 18 °C, at 300 r.p.m. Proteins were refolded and purified from inclusion bodies using a single freezing-thawing method^92^. In brief, cells were harvested and lysed by sonication in 40 ml PBS (pH=8.0) supplemented with 1 mM PMSF and1×EDTA-free protease inhibitor cocktail (#11873580001, Roche) followed by centrifugation at 4 °C, 30 min, 21,000*xg*. Inclusion bodies were washed three times in 40 ml buffer containing 20 mM Tris (pH 8.0), 300 mM NaCl, 1 mM EDTA, 1% Triton X-100, and 1M urea, and centrifuged at 4 °C, 20 min, 12,000*xg*. Finally, the pellet was resuspended in PBS supplemented with 2M urea and stored at -20 °C for 24h. Frozen samples were thawed slowly on ice and centrifuged at 4 °C, 20 min, 12,000*xg*. Supernatant containing the resolubilized recombinant FusB was collected and dialyzed over PBS supplemented with 0.65 M urea at 4 °C for 24h, followed by a second dialysis in PBS at 4 °C for 16h. After centrifugation at 4 °C, 10 min, 5,000 *xg*, supernatant was collected, filter sterilized and FusB was concentrated using an Amicon Ultra centrifugal device with 30 kDa molecular weight cut-off (#UFC9030, Merck Millipore) at 4 °C, 60 min, 4,000*xg*. Samples were flash-frozen in liquid nitrogen and stored at -80 °C before use. Concentration and integrity of the ∼77 kDa FusionB was assessed by SDS-PAGE analysis followed by Coomassie Brilliant Blue-staining or western blotting (Figure S17) using anti-cdtB-B or anti-aip56-B monoclonal antibodies, respectively^38^.

### Nuclease assays

Supercoiled pGEM-7zf(+) plasmid was isolated from overnight cultures using the Monarch Plasmid Miniprep Kit (NEB) at 4°C. Purified plasmid was added to a total concentration of 250ng/reaction in reaction buffer (25mM HEPES, 5mM CaCl_2_, 5mM MgCl_2_, 1mM DTT). 0.1-1µg purified protein (either *Da*CdtB or FusionB) was added to each reaction to a total reaction volume of 20µl. Negative controls for all reaction conditions consisted of plasmid, reaction buffer ± reducing agent without protein. Reactions were incubated for 1h at 28°C. Reactions were deactivated with 0.5µl Proteinase K (NEB) + 0.5% SDS at 55°C for 10 minutes. A linearized control was generated by digesting the plasmid with EcoRI-HF (NEB) in rCutSmart buffer (NEB) at 37°C for 1h, followed by heat inactivation at 65°C for 20 min. The full 20ul reactions were suspended in no-SDS purple loading dye (NEB) and run on a 0.8% TBE agarose gel with SYBR Safe (Invitrogen) at 75V for 2h. Gels were visualized on a Gel Doc XR+ visualization system (BioRad).

### Survival assays

Five balanced UAS or *Actin-GAL4* virgin females and 2 balanced UAS or *Actin-GAL4* males were set in crosses. The parental generation was flipped onto new media after 1 week of laying to allow for accurate scoring of F1 offspring. Offspring were counted and phenotyped for the balancer for one week following the emergence of the first F1 flies. The deviation from Mendelian expectation was calculated as:

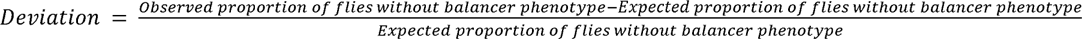

For crosses with UAS on chromosome 2, the expected Mendelian proportion of flies without the balancer phenotype was 0.33, due to the homozygous lethal nature of CyO/CyO. For crosses with UAS on chromosome 3, the expected Mendelian proportion of flies without the balancer phenotype was 0.25.

### Developmental time course

Ten homozygous UAS-*fusionB* females were crossed to 5 Actin-GAL4/CyO, GFP male flies and allowed to lay in chambers outfitted with grape agar plates supplemented with live yeast paste. Fresh plates were switched every four hours to synchronize larval development. First or second instar larvae were sorted under a Nightsea GFP scope to separate balancer (GFP+) and non-balancer (GFP-) siblings, which were then moved into vials containing standard molasses food to complete development. Vials were monitored each day for pupal development. Time to pupariation for each larvae and the proportion of flies reaching pupariation were recorded for each cross. Staged pupae were photographed at 24 hour intervals using a Keyence VHX-5000 digital microscope, using the 20-200× lens at 150×.

### Generation of yeast strains

Inserts were cloned into the pENTR vector using the Gateway pENTR/D-TOPO system (Invitrogen). Transformants were sequence verified before isolation with the Monarch Plasmid Miniprep Kit (NEB). These vectors were recombined into the pAG425-ccdB (Addgene #14153)^93^ plasmid using the Gateway LR Clonase II Enzyme Mix (Invitrogen) to generate galactose-inducible vectors for yeast expression. Resulting clones were sequence verified, then transformed into *Saccharomyces cerevisiae* BY4743 using the LiAc/SS carrier DNA/PEG method^94^. Yeast transformants were screened using the NaOH DNA preparation^95^ followed by PCR amplification of the insert, then were sequence verified before use in experiments. At all stages, Sanger sequencing was used to verify insert sequences <1kb and Plasmidsaurus/Primordium sequencing was used to verify insert sequences >1kb.

### Yeast toxicity assay

Single colonies were picked and grown in liquid CSM-Leu + 2% glucose overnight. Cultures with an OD_600_ < 1 were excluded from the analysis. Cultures were then spun down, washed in water, diluted to an OD_600_ 0.2, then serially diluted ten-fold to a final dilution of 10^-4^. Serial dilutions were plated on CSM-Leu + 2% glucose (non-inducing) or CSM-Leu + 2% galactose (inducing) in 5µl volumes. Spot plates were grown at 30°C for 72 hours. Growth was measured by counting colony forming units (CFUs) as well as spot density. Spot density was measured using the protocol described in Petropavlovskiy et al. (2020)^96^, modified to be compatible with light colored backgrounds. Briefly, background subtraction steps were conducted in ImageJ as described in [96]. Mean grey values were measured for spots at the 10^-1^ dilution, as this was the lowest dilution with consistent visible differences across experiments. Mean grey values for each spot were subtracted from the background to generate a true mean grey value for each spot. Mean grey values were then normalized to the mean grey value of the empty vector control in each image for comparison.

### Replica plating experiment

Yeast cultures were grown and diluted as described in the toxicity assay. 50µl of a 10^-2^ dilution of each strain was plated onto CSM-Leu + 2% galactose. Yeast were allowed to grow for 48h (empty vector and truncated *fusion* vectors) or 72h (*fusion* vectors) before replica plating onto YPG and YPD plates. The difference in incubation times before replicating was to control for colony size prior to replicating to minimize bias in initial inoculum sizes. YPG and YPD plates were scanned for growth at 24h and 48h post replica plating. Colonies were counted at the 48h timepoint and the fraction of growth on each substrate was calculated by dividing the number of colonies on each plate by the number of colonies on the CSM-Leu parent plate.

## Key Resources

| REAGENT or RESOURCE | SOURCE | IDENTIFIER |
| --- | --- | --- |
| **Antibodies** |  |  |
| anti-AIP56 2H5 | Verster, Cinege et al. 2023<sup>38</sup> | N/A |
| Anti-CdtB 3G9 | Verster, Cinege et al. 2023<sup>38</sup> | N/A |
| anti-lamellocyte L1 | Kurucz et al. 2007<sup>90</sup> | N/A |
| Alexa Fluor 488 goat anti-mouse IgG | Invitrogen | Catalog #A32723 |
| Polyclonal Goat Anti-Mouse Immunoglobulins/horseradish peroxidase | Dako/Agilent | Part Number: P044701-5 |
| **Bacterial and virus strains** |  |  |
| SixPack *Escherichia coli* | Lipinski et al. (2018)<sup>91</sup> | N/A |
| **Chemicals, peptides, and recombinant proteins** |  |  |
| *D. ananassae* CdtB | Verster et al. 2019<sup>36</sup> | N/A |
| *D. ananassae* FusionB | This paper | N/A |
| EcoRI-HF | New England Biolabs | Catalog #R3101S |
| NotI-HF | New England Biolabs | Catalog #R3189S |
| **Critical commercial assays** |  |  |
| Q5 High-Fidelity DNA Polymerase | New England Biolabs | Catalog #M0491S |
| Protoscript II First Strand cDNA Synthesis Kit | New England Biolabs | Catalog #E6560S |
| NEBuilder HiFi DNA Assembly Master Mix | New England Biolabs | Catalog #E2621S |
| pENTR/D-TOPO Cloning Kit | Invitrogen | Catalog # K240020SP |
| Gateway LR Clonase II Enzyme Mix | Invitrogen | Catalog #11791020 |
| Monarch Plasmid Miniprep Kit | New England Biolabs | Catalog #T1010S |
| **Experimental models: Organisms/strains** |  |  |
| *Saccharomyces cerevisiae* strain BY4743 | Nicholas Ingolia | ATCC® Catalog No.: 201390 |
| *Drosophila ananassae* | UC San Diego Species Stock Center | 14024-0371.13 |
| *Drosophila melanogaster* UAS-*H. defensa cdtB* (attp40) | This study | N/A |
| *D. melanogaster* UAS-*D. ananassae cdtB* (attp40) | This study | N/A |
| *D. melanogaster* UAS-*D. ananassae fusionA* (attp40) | This study | N/A |
| *D. melanogaster* UAS-*D. ananassae fusionA* (attp2) | This study | N/A |
| *D. melanogaster* UAS-*D. ananassae fusionB* (attp40) | This study | N/A |
| *D. melanogaster* UAS-*D. ananassae fusion* (attp2) | This study | N/A |
| *D. melanogaster* UAS-*gfp* | Bloomington Stock Center | RRID:BDSC_5431 |
| *D. melanogaster* Actin-GAL4 | Bloomington Stock Center | RRID:BDSC_4414 |
| *D. melanogaster* Actin-GAL4/CyO, GFP | Iswar Hariharan | N/A |
| *D. melanogaster* Cg-GAL4 | Bloomington Stock Center | RRID:BDSC_7011 |
| *D. melanogaster* He-GAL4 | Bloomington Stock Center | RRID:BDSC_8699 |
| *Leptopilina boulardi* Lb17 | Todd Schlenke | N/A |
| *Leptopilina boulardi* G486 | Todd Schlenke | N/A |
| *Leptopilina heterotoma* Lh14 | Todd Schlenke | N/A |
| **Oligonucleotides** |  |  |
| Primers for cloning and strain validation, see table S6. | This study | N/A |
| **Recombinant DNA** |  |  |
| For details on gene blocks synthesized for this study, see table S5. | This study | N/A |
| Plasmid: pUAST::attB | Bischof et al. (2007)<sup>89</sup> |  |
| Plasmid: pAG425-ccdB | Alberti et al. (2007)<sup>93</sup> | Addgene #14153 |
| Plasmid: pGEM-7zf(+) | Promega | Catalog #P2251 |
| **Software and algorithms** |  |  |
| BLAST | Altschul et al. 1997<sup>67</sup> | https://blast.ncbi.nlm.nih.gov/Blast.cgi |
| cd-hit v 4.8.1 | Li & Godzik (2006)<sup>68</sup> | https://sites.google.com/view/cd-hit |
| MUSCLE v 5.1.linux64 | Edgar (2004)<sup>69</sup> | https://www.drive5.com/muscle/ |
| AliView v 1.28 | Larsson (2014)<sup>70</sup> | https://ormbunkar.se/aliview/ |
| Clipkit webserver | Steenwyk et al. <sup>71</sup> |  |
| IQTree Webserver | Nguyen et al. (2015)<sup>72</sup> | http://iqtree.cibiv.univie.ac.at/ |
| ggtree v 3.10.1 | Yu et al. (2017)<sup>73</sup> | https://guangchuangyu.github.io/software/ggtree/ |
| ggtreeExtra v 1.12.0 | Xu et al. (2021)<sup>74</sup> | https://github.com/YuLab-SMU/ggtreeExtra |
| phytools v 2.1.1 | Revell (2024)<sup>75</sup> |  |
| ape v 5.7.1 | Paradis & Schliep (2019)<sup>76</sup> | https://github.com/emmanuelparadis/ape |
| Progressive Cactus v.2.6.0 | Armstrong et al. (2020)<sup>80</sup> | https://github.com/ComparativeGenomicsToolkit/cactus |
| Comparative Annotation Toolkit software v2.2.1 | Fiddes et al. (2018)<sup>81</sup> | https://github.com/ComparativeGenomicsToolkit/Comparative-Annotation-Toolkit |
| Geneious Prime 2023.1.2 | Dotmatics | https://www.geneious.com/ |
| RStudio 2021.09.0 | Posit | https://posit.co/download/rstudio-desktop/ |
| Mafft v 7.490 | Katoh & Standley (2013)<sup>83</sup> | https://mafft.cbrc.jp/alignment/software/ |
| Berkeley Drosophila Genome Project Splice Site Prediction by Neural Network | Reese et al. (1997)<sup>84</sup> | https://www.fruitfly.org/seq_tools/splice.html |
| AlphaFold Multimer | Evans et al. (2022)<sup>85</sup> | https://cosmic-cryoem.org/tools/alphafoldmultimer/ |
| ColabFold v 1.5.5 | Mirdita et al. (2022)<sup>86</sup> | https://colab.research.google.com/github/sokrypton/ColabFold/blob/main/batch/AlphaFold2_batch.ipynb |
| ChimeraX v 1.7 | Meng et al. (2023)<sup>88</sup> | https://www.cgl.ucsf.edu/chimerax/ |
| ImageJ v 2.14.0/1.54f | National Institutes of Health | https://imagej.net/ij/index.html |
| **Other** |  |  |
| For accession numbers for genetic data used for annotating HGT loci, see table S4. | This study | N/A |

## Acknowledgments

We thank Genetivision Corporation for establishing all new fly lines used in this study. We thank L. Brennan and E. Ünal for discussions on experimental design for the biochemical and yeast experiments, respectively. We thank J. Massey for helpful feedback on project development and early drafts of the manuscript. We thank J. Peláez for providing vector cartoons of fly larvae.

R.L.T. was funded by the National Institutes of Health Genetic Dissection of Cells and Organisms Training Grant (award #5T32GM132022-03) and the National Science Foundation Graduate Research Fellowship. This project was supported by funding from the National Institute of General Medical Sciences of the National Institutes of Health (award no. R35GM119816) to N.K.W., the NKFI K135877 grant from the Hungarian National Science Foundation to I.A., and the National Laboratory for Biotechnology (2022-2.1.1-NL-2022-00008) and the Hungarian Academy of Sciences (Lendület Program Grant (LP2017-7/2017)) to Z.L..

## Author contributions

R.L.T. and N.K.W. conceived the project. R.L.T. designed the experiments with contributions from G.C., K.I.V., Z.L., I.A., and N.K.W. B.Y.K. performed the initial genome annotation analysis and R.L.T. manually adjusted annotations. R.L.T. designed all fly transgenic lines. J.A.T. and R.L.T. performed fly overexpression and parasitization experiments. G.C. performed indirect immunofluorescence and western blot experiments, with help from E.A., L.B.M., J.A.T., and S.L.B. J.H.H. and R.L.T. performed yeast expression experiments. Z.L. performed protein purification of recombinant FusionB. I.A. generated monoclonal antibodies against *D. ananassae* CdtB and Fusion genes and performed parasitization and indirect immunofluorescence assays to survey hemocyte differentiation. R.L.T. and N.K.W. drafted the manuscript, and all authors provided comments and contributed to editing the manuscript.

## List of Supplementary Materials

Document S1. Figs. S1-S17 and Tables S1-S3

Document S2. Excel file with Tables S4-S6 which were too large for PDF, related to STAR Methods.

