## Supplemental File1 (Figs. S1-S17, Tables S1-3) for "Retracing the horizontal transfer of a novel innate immune factor in *Drosophila*"

**Supplementary material for: Retracing the horizontal transfer of a novel innate immune factor in *Drosophila***

**Materials & Methods**

**Supplementary Figures 1-17**

**Supplementary Tables 1-8**

**Materials & Methods**

**Protein phylogenies**

Deviation = Observed proportion of flies without balancer phenotypeExpected proportion of flies without balancer phenotype

For crosses with UAS on chromosome 2, the expected Mendelian proportion of flies without the balancer phenotype was 0.33, due to the homozygous lethal nature of CyO/CyO. For crosses with UAS on chromosome 3, the expected Mendelian proportion of flies without the balancer phenotype was 0.25.

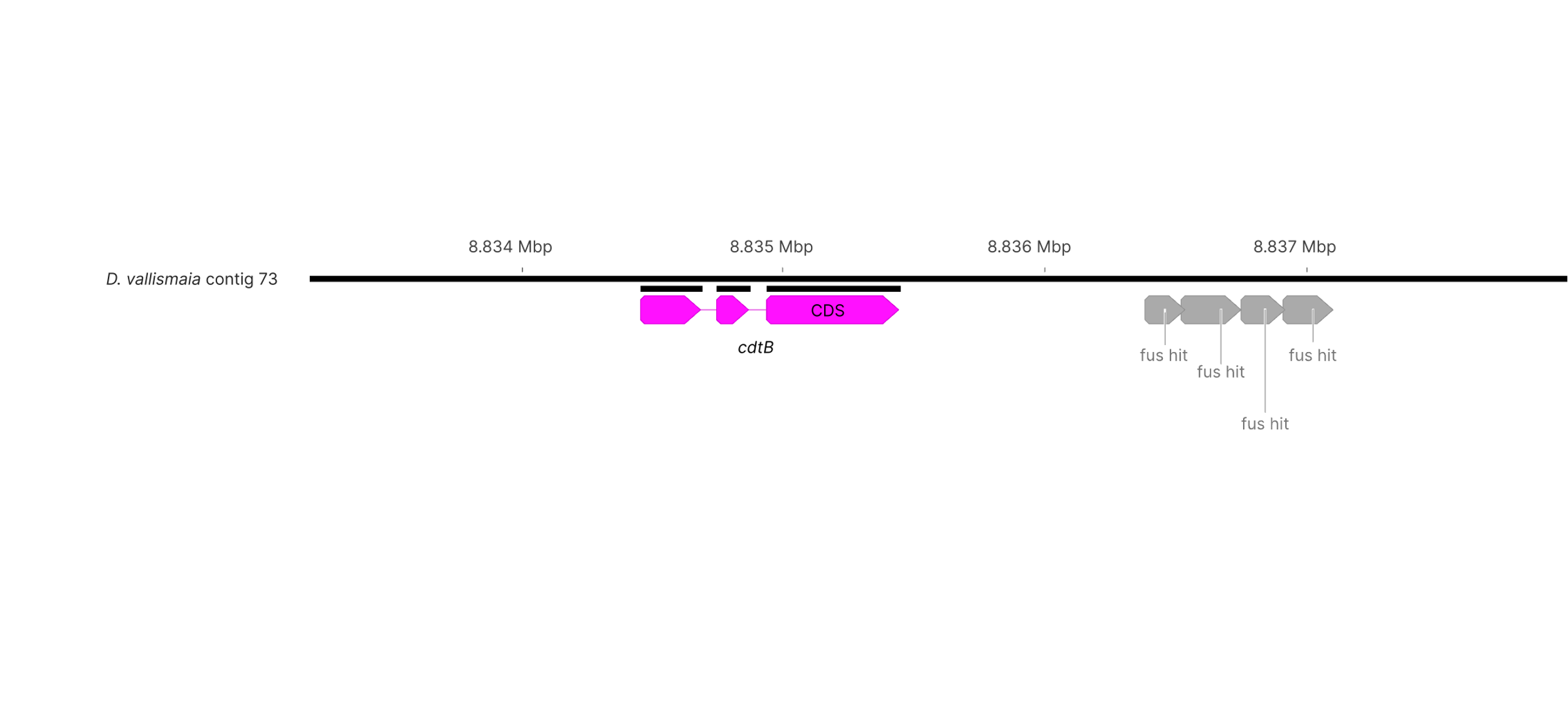
**Figure S1. *D. vallismaia* genome contains remnants of an ancestral *fusion* gene.** *D. bipectinata* Fusion protein was queried in a tblastn search against the *D. vallismaia* genome in Geneious Prime 2023.1.2. The coordinates of significant hits were mapped.

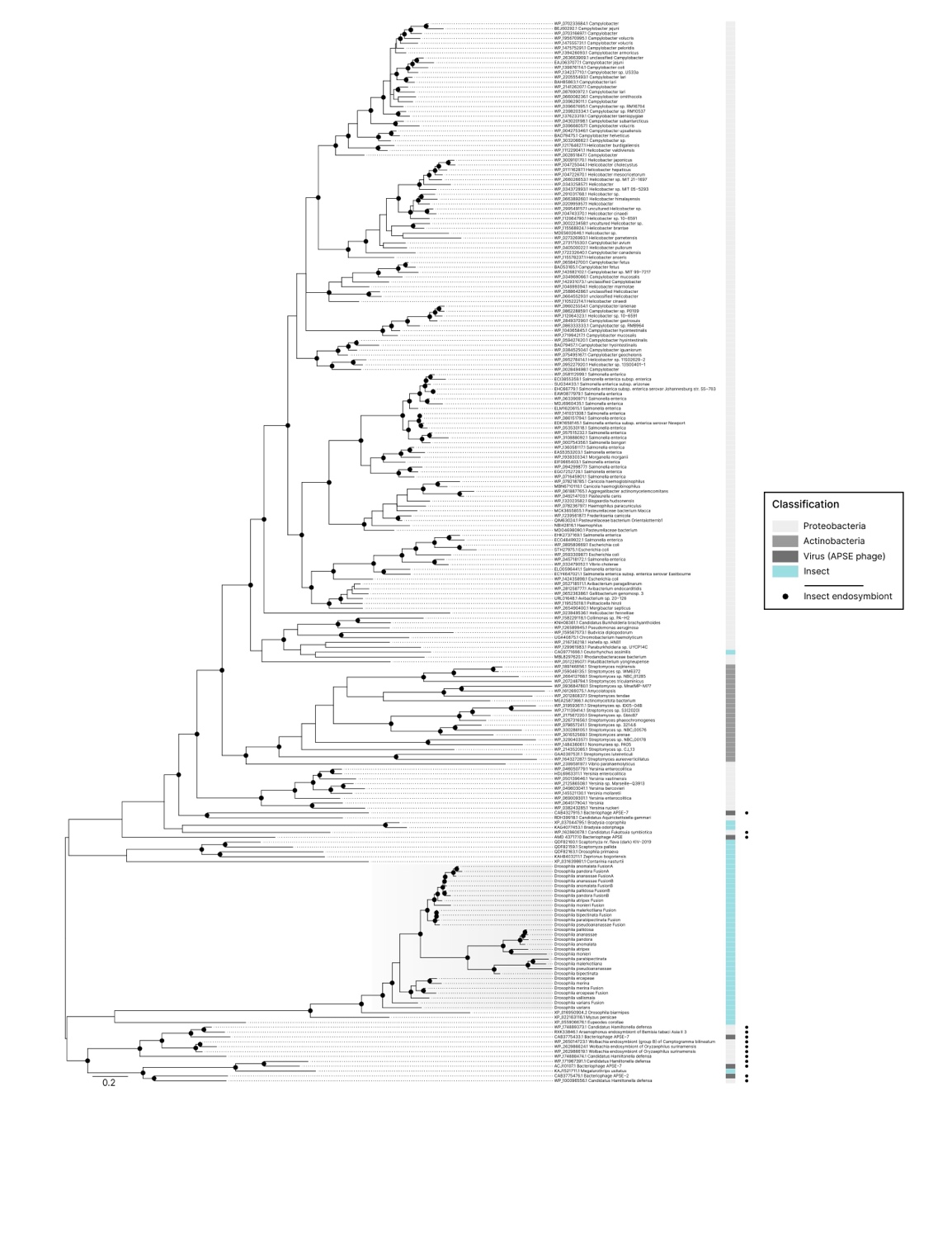

A

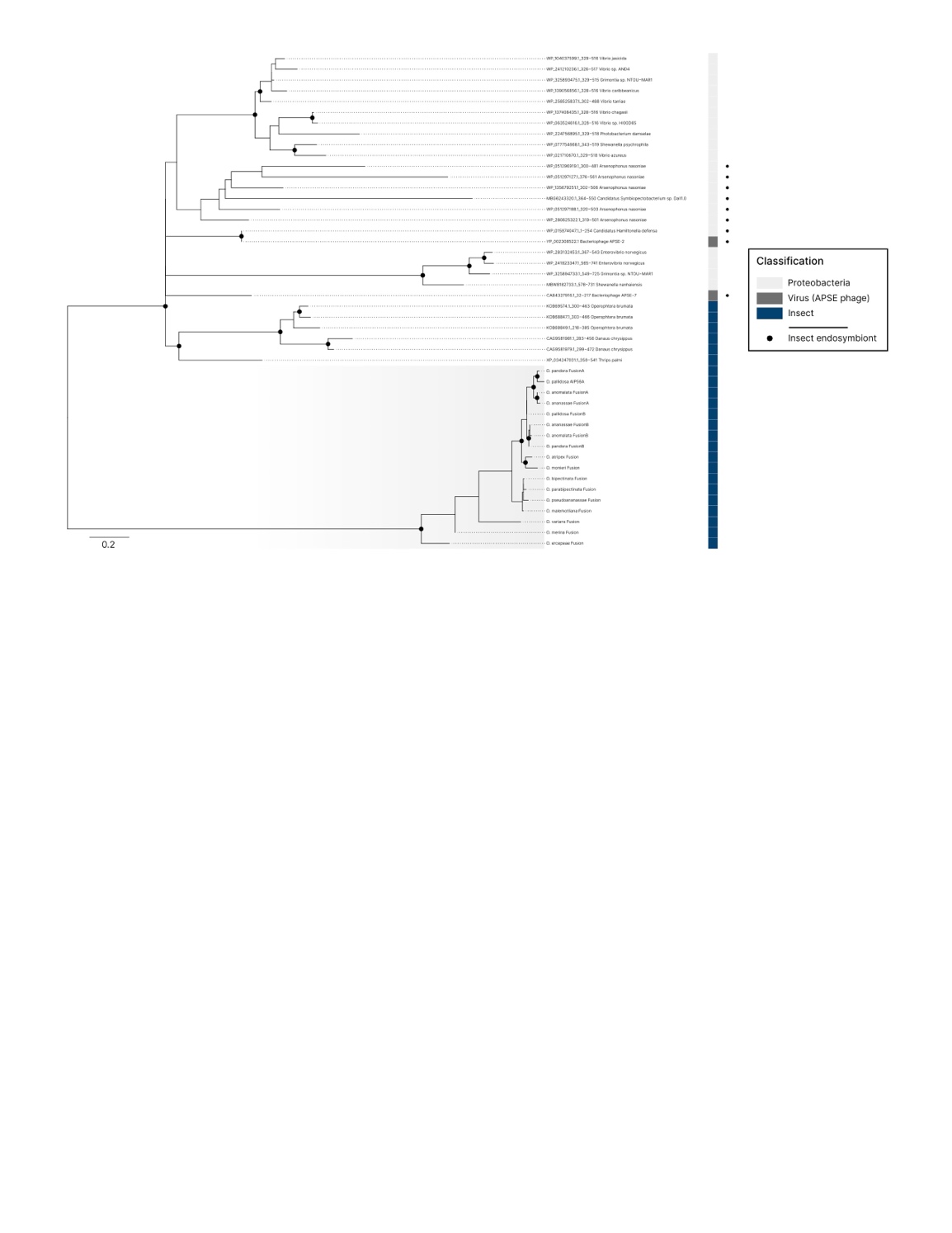

B

**Figure S2. CdtB and AIP56 protein trees.** CdtB (A) and AIP56 (B) protein trees indicate *ananassae* group CdtB and AIP56 proteins are most related to homologs deriving from insect endosymbionts. Both maximum likelihood gene trees were generated in IQTree using the best fit model (CdtB: VT+F+G4; AIP56: WAG+F+I+G4). Trees are midpoint rooted. Nodes with <50% bootstrap support are collapsed into polytomies, and nodes with >90% bootstrap support are labelled. *ananassae* subgroup homologs are highlighted. Scale bar = substitutions/site.

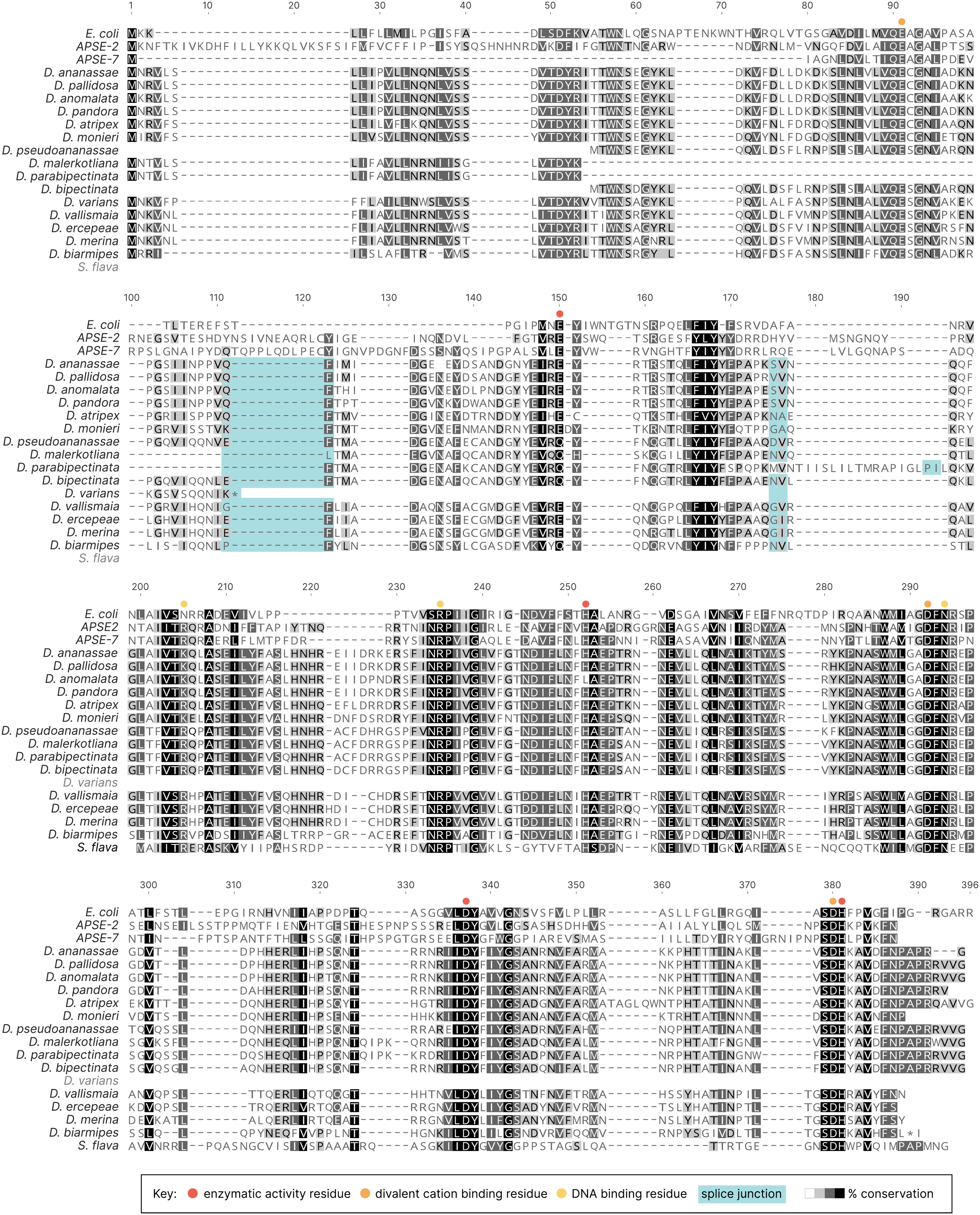
**Figure S3. CdtB protein alignment.** Alignment of *ananassae* group CdtB proteins indicates high conservation of CdtB functional residues^43^ and splice junctions. CdtB protein alignment was generated using MAFFT v7.490 as implemented in Geneious Prime 2023.1.2. Residue conservation is scored using the BLOSUM62 matrix.

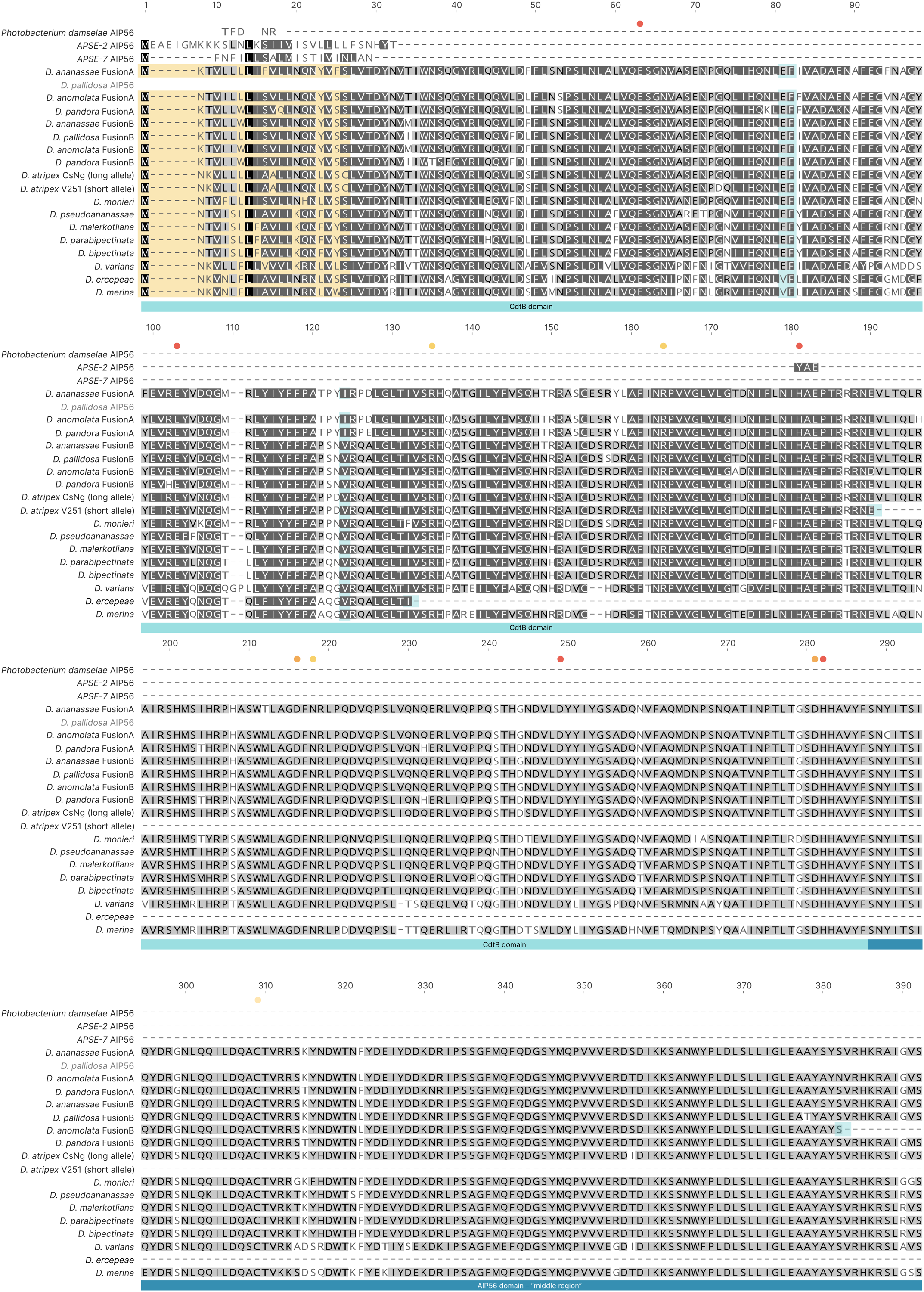
­­

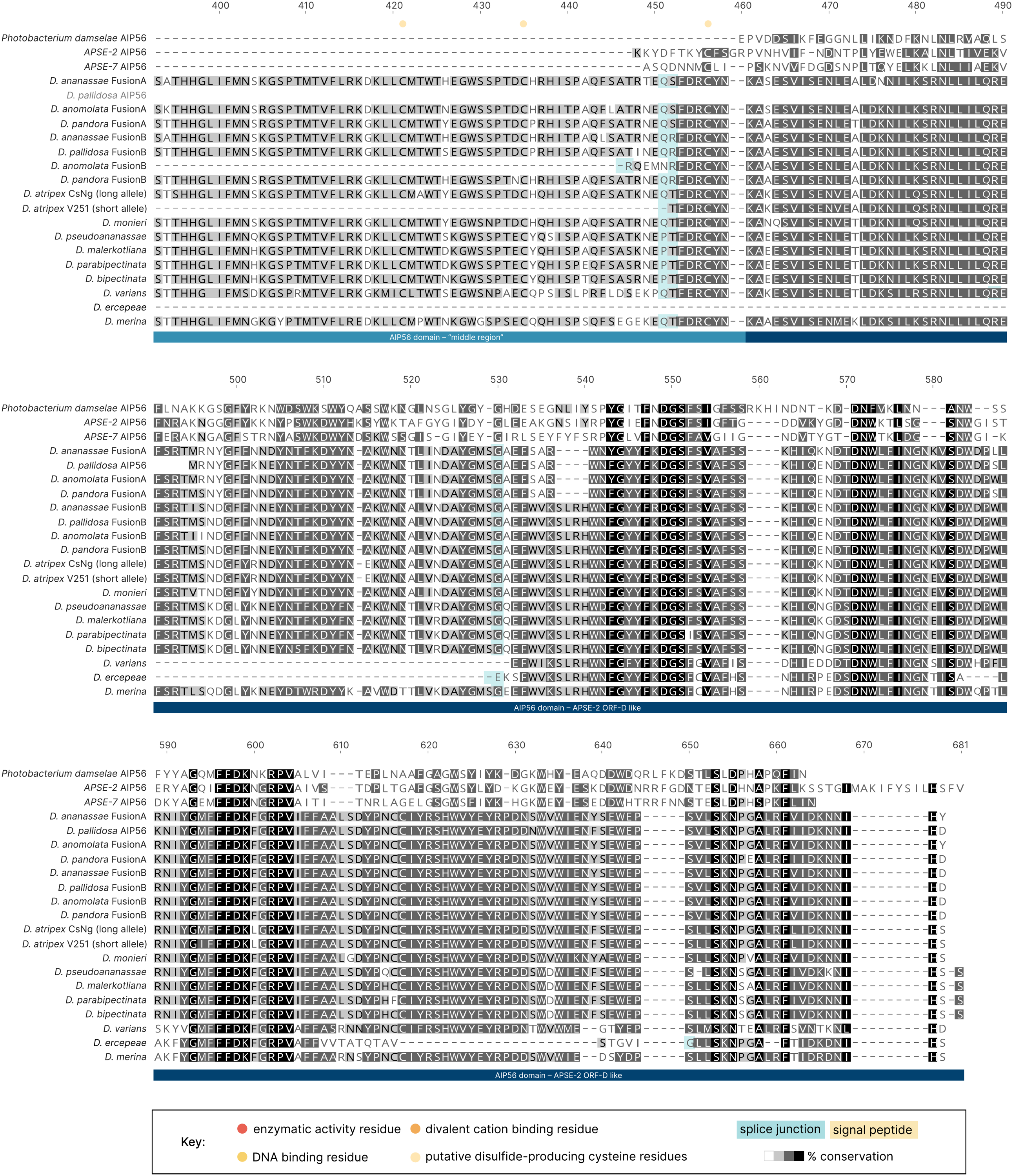
**Figure S4. Fusion protein alignment.** Alignment of *ananassae* group Fusion proteins indicates high conservation of CdtB functional residues^43^ and splice junctions among the Fusion genes. Fusion protein alignment was generated using MAFFT v7.490 with the BLOSUM62 matrix as implemented in Geneious Prime 2023.1.2. Residue conservation is scored using the BLOSUM62 matrix.

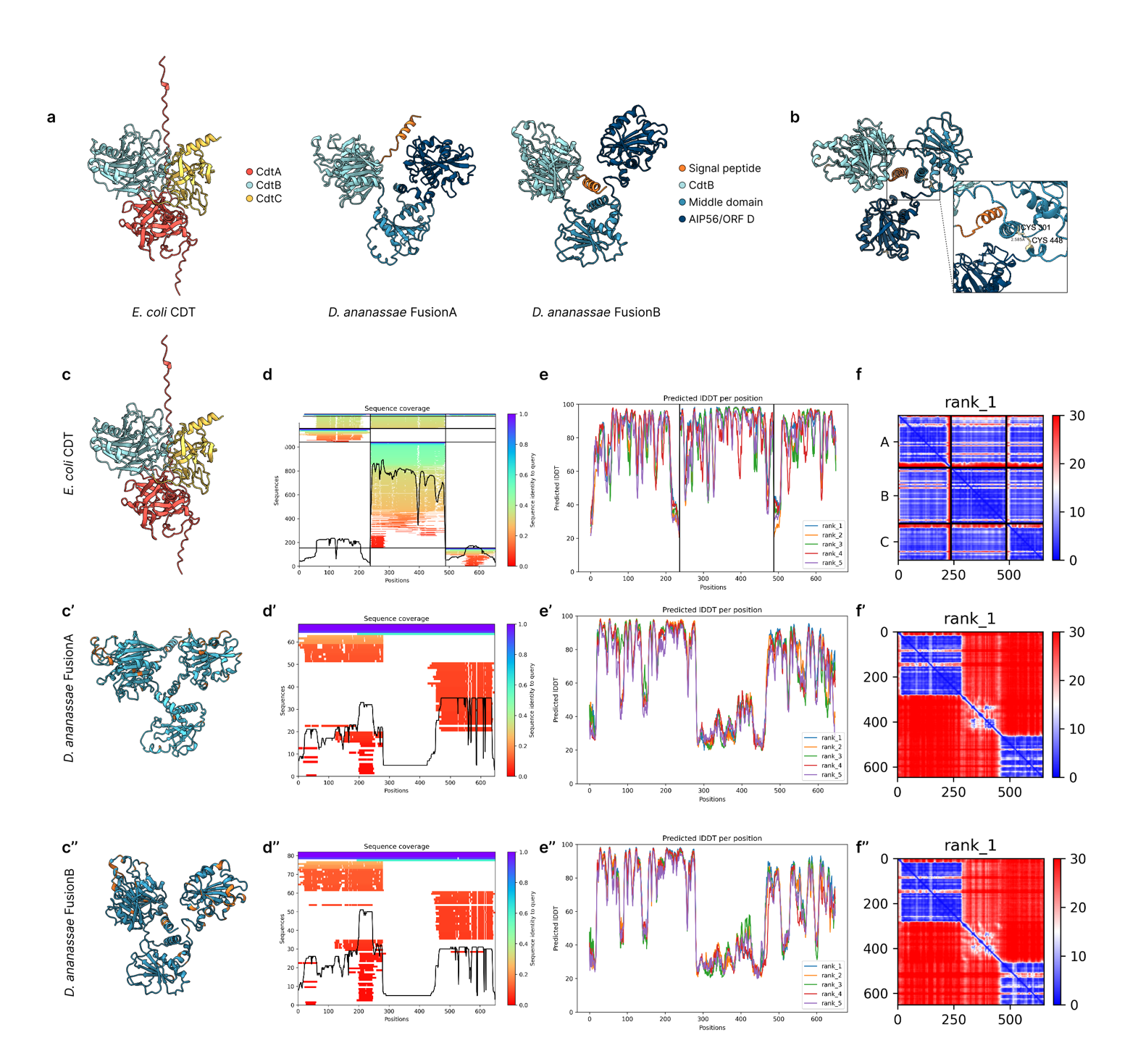
**Figure S5. AlphaFold output for FusionA and FusionB proteins and structural similarity to CDT holotoxins. (**A) AlphaFold predictions for *E. coli* CDT (left) and *D. ananassae* FusionA and FusionB holotoxins. Structures are colored by domain identity. (B) Putative disulfide bond site between FusionB C301 and C448. Alphafold structural prediction for FusionB protein places sulfur residues on C301 and C448 within 2.585Å (inset). Typical disulfide bonds are 2.05Å. (C-C’’) Structure predictions for *E. coli* CDT (C), *D. ananassae* FusionA (C’), and *D. ananassae* FusionB C’’). Orange residues indicate nonsynonymous sites between *D. ananassae* FusionA and FusionB. (D-D’’) Sequence coverage plots for Alphafold predictions. (E-E’’) Per-residue confidence metric (pLDDT) output for each structure. (F-F’’) Predicted aligned error (PAE) for each structure.

**Figure S6. *D. atripex* genotyping**
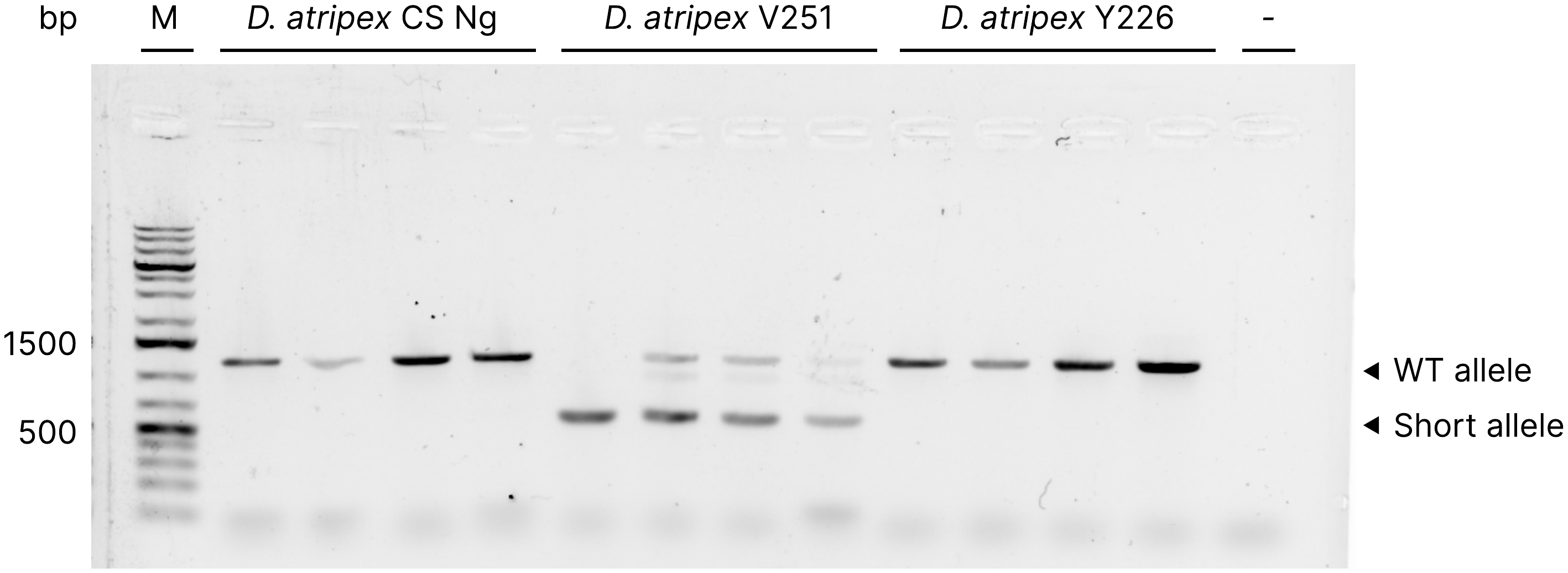
**indicates segregating structural polymorphism in strain V251.** Each lane represents gDNA isolated from a single individual. - = no DNA control.

**Figure S7. Hemocytes proliferate normally in *fusionB*-expressing flies.** Indirect immunofluorescence performed on naive (A-B) or *L. boulardi-*infected (C-D) *D. melanogaster* hemocytes, stained for either Fusion expression (AIP56, panels A and C) or a lamellocyte-specific marker (Lamellocytes, panels B and D). (A) FusionB is detected in *He-Gal4 > fusB* hemocytes, but not in hemocytes from either parental strain. (B) Lamellocytes fail to form in the absence of parasitoid challenge in all three *D. melanogaster* genotypes. (C) FusionB is detected only in *He-Gal4 > fusB* hemocytes. Lamellocytes are able to differentiate in all three strains and are indicated with white triangles. (D) *He-Gal4 > fusB* and its parent strains form lamellocytes following parasitoid challenge. Scale bar = 50µm.

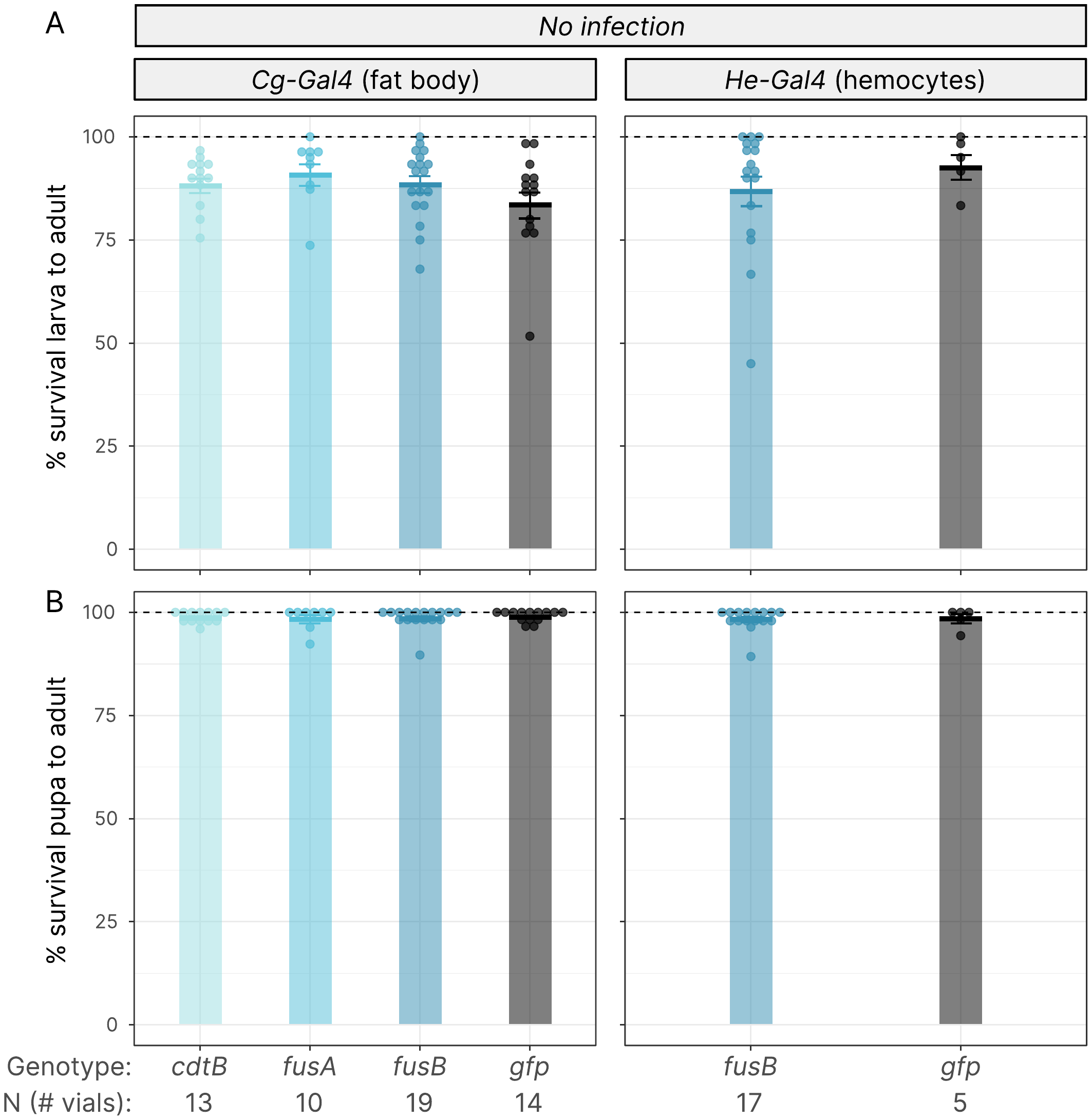
**Figure S8. Transgene expression alone does not impair fly survival. (**A) Larval to adult survival proportions for all crosses. (B) Pupa to adult survival proportions for all crosses. Once flies reach pupal stage, they almost always complete development to adult. Crossbars represent mean ± standard error.

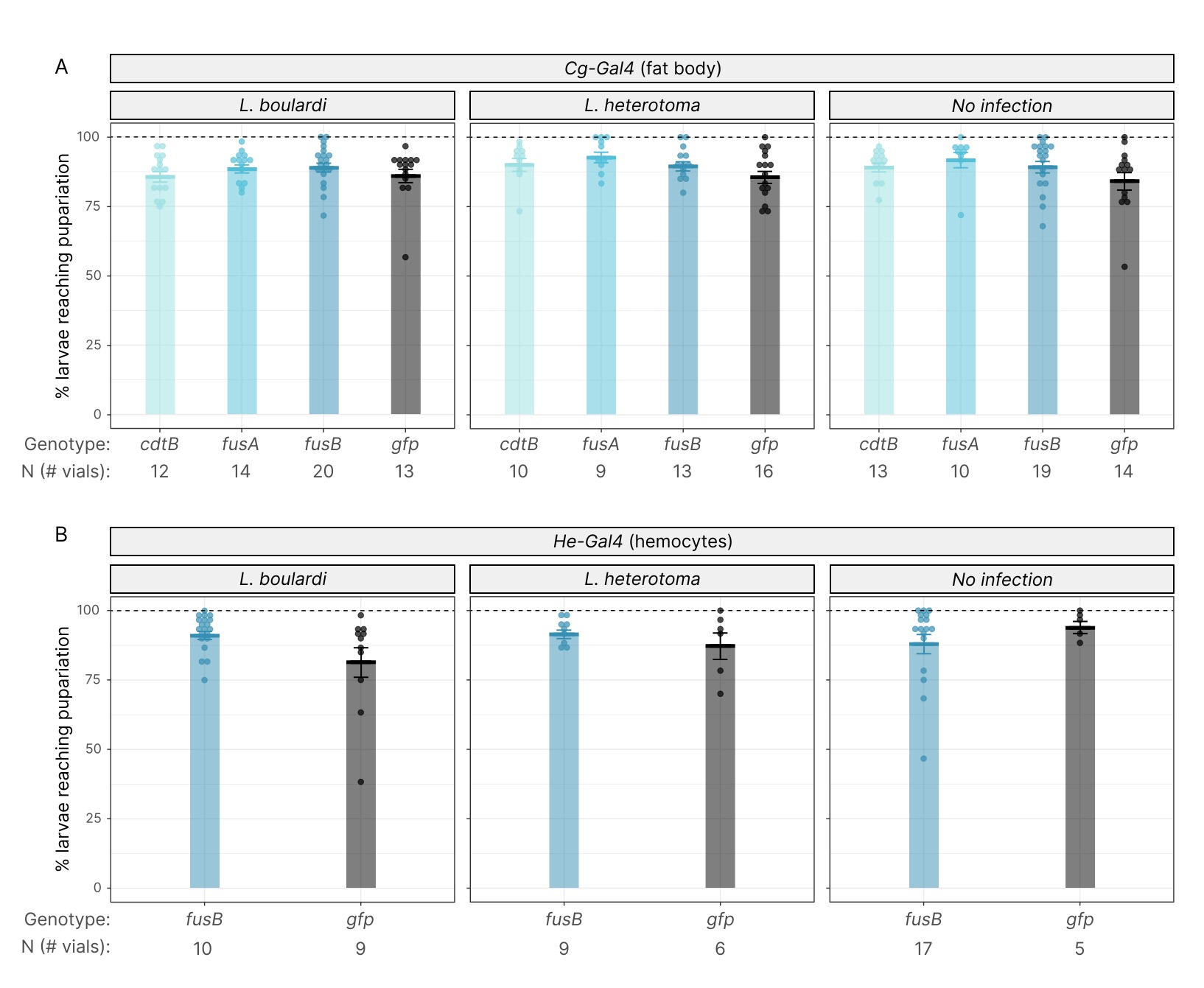
**Figure S9. Pupariation rates for all genotypes across all infection schemes.** In both *Cg-Gal4* (A) and *He-Gal4* (B) crosses, all genotypes reached pupariation at similar rates, regardless of infection status. Crossbars indicate mean ± standard error.

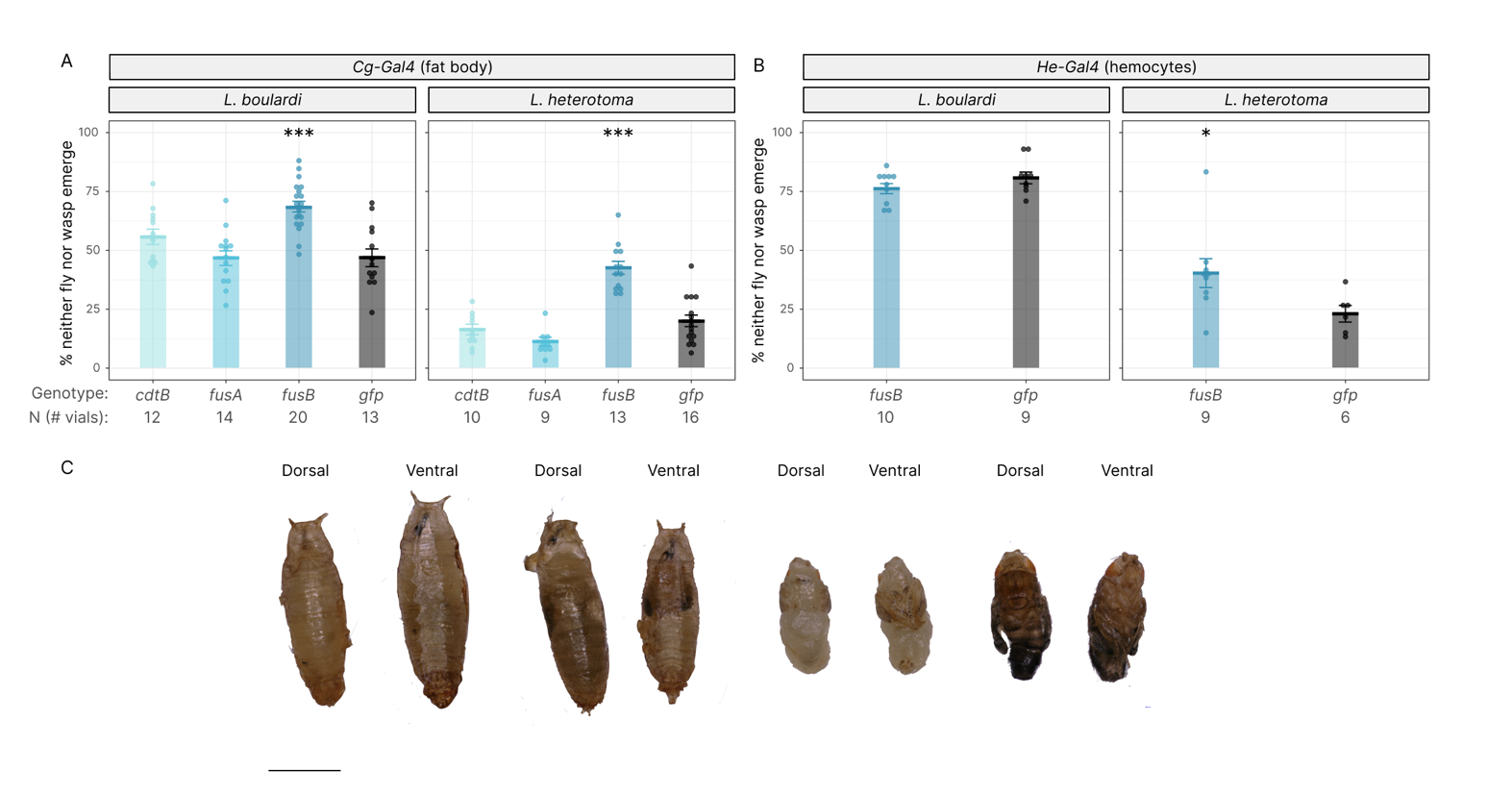
**Figure S10. *fusionB*-expressing flies have higher incidence of stalemates with wasps.** (A-B) Proportion of larvae from which neither an adult fly or an adult wasp complete development in *Cg-Gal4* (A) or *He-Gal4* (B) crosses. * = p < 0.05, *** p < 0.001 per pairwise Wilcoxin rank sum tests. Crossbars represent mean ± standard error. (C) *Cg-Gal4 > fusB* pupae that arrested at various stages following infection with *L. boulardi.* Pupae are ordered from least advanced development to most advanced. Less advanced pupae were kept in pupal case for imaging, and the dorsal and ventral images are two different representative pupae that arrested around the same stage. “Advanced” pupae were dissected from pupal cases and imaged from their dorsal and ventral sides. Scale bar = 1 mm.

**Figure S11. *Cg-Gal4* > *fusB* phenocopies native *D. ananassae fusion* expression patterns.** General morphology of Fusion protein signals are highly similar between *D. ananassae* and *D. melanogaster Cg-Gal4 > fusB*. *D. ananassae* samples were prepared and imaged separately from the *D. melanogaster* samples, so signal intensity should be interpreted with caution. Scale bar = 50µm.

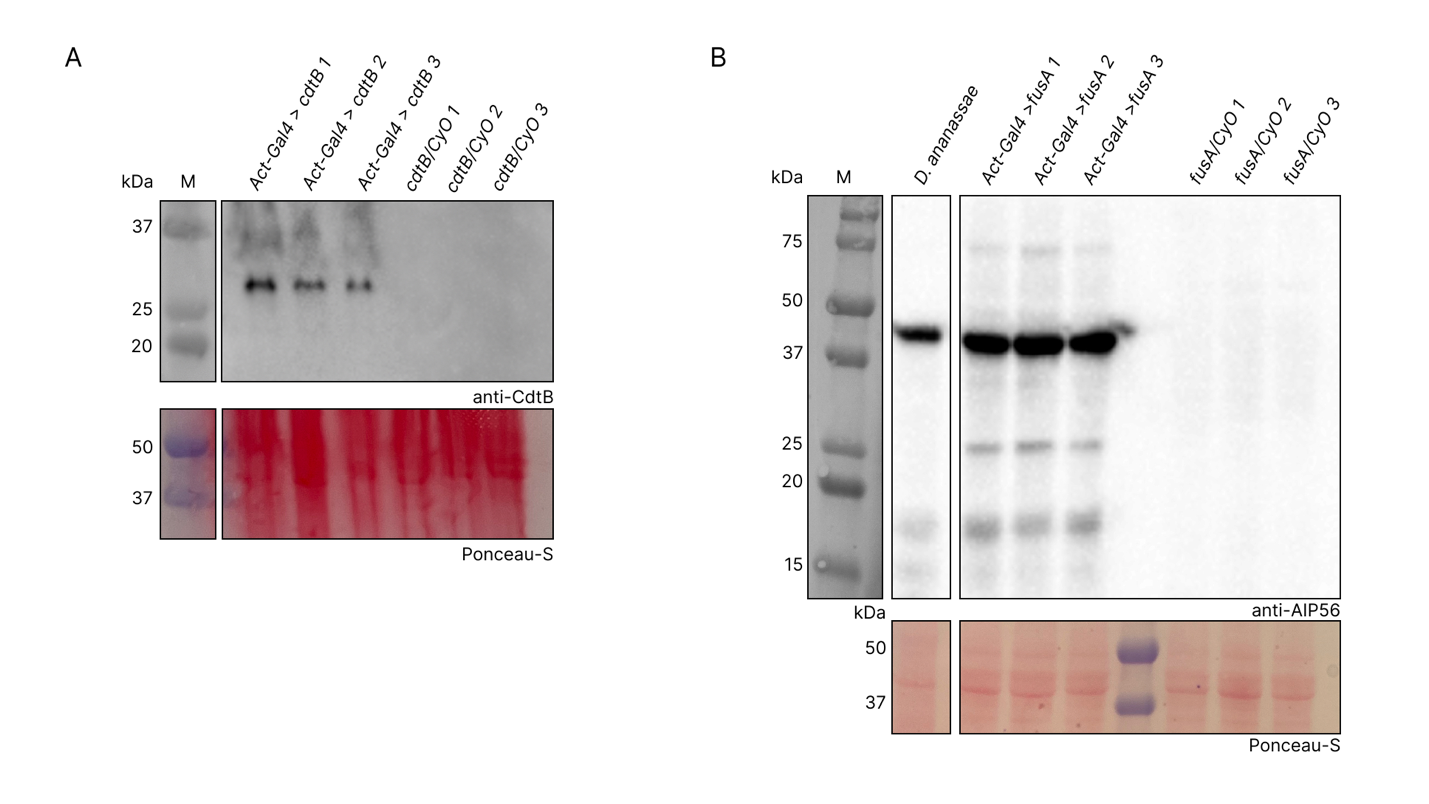
**Figure S12. CdtB and FusionA are expressed under *Actin-Gal4.*** Western blots targeting *D. ananassae* CdtB (A) and the AIP56 domain of FusionA (B) demonstrate the proteins are translated in flies that do not have the balancer phenotype. Flies with the balancer phenotype (CyO) do not express these genes. Each lane represents samples taken from siblings from 3 independent crosses.

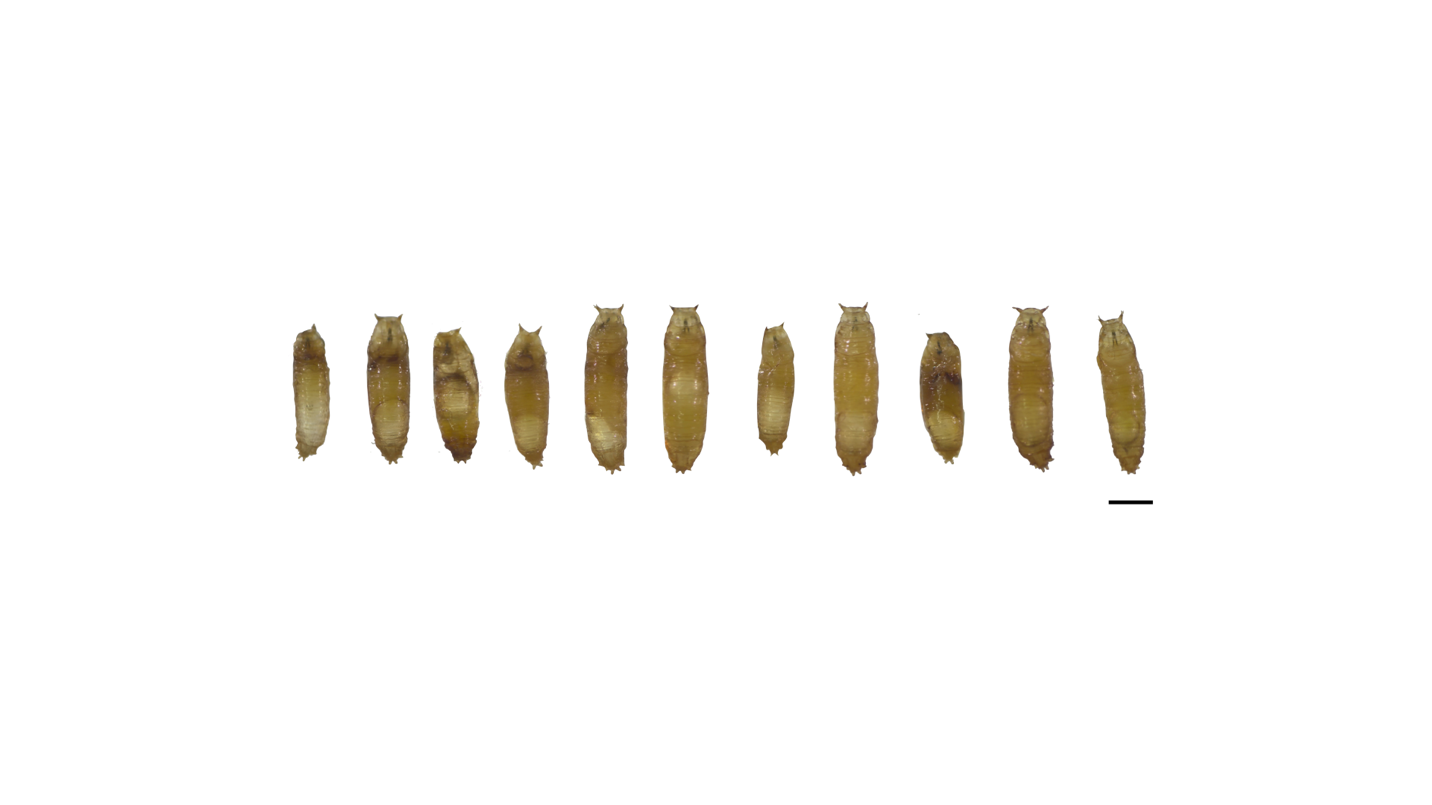
**Figure S13. Representative images of *fusB/Act-Gal4*** **pupae.** Pupae staged at 96h after pupariation. Scale bar = 1mm.

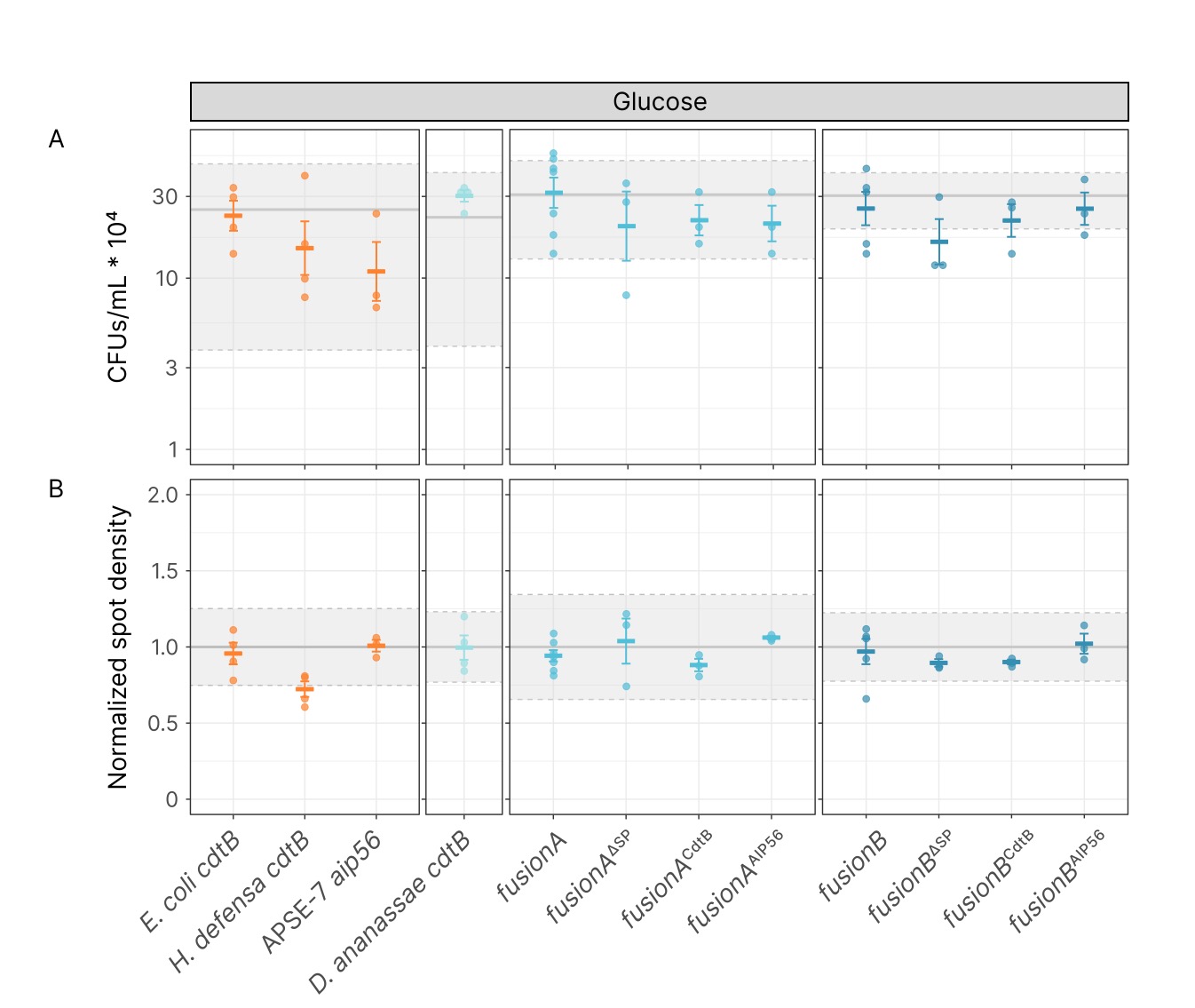
**Figure S14. All strains grow comparably on non-inducing media.** CFU counts (A) and normalized spot densities (B) for all strains grown on glucose (non-inducing) media. Crossbars indicate mean ± standard error. Grey boundaries indicate the mean (solid line) and 95% confidence interval (grey rectangle) of CFU counts for empty vector controls used in experiments with each strain.

**Figure S15. Drosophilid CdtB is generally non-toxic to yeast.** A) Representative image of spot assays on CSM-Leu media containing either 2% glucose (non-inducing, left) or 2% galactose (inducing, right). B) CFU counts for each strain on galactose (left) and glucose (right) media. Cross bars indicate mean ± standard error. Grey boundaries indicate the mean (solid line) and 95% confidence interval (grey rectangle) of CFU counts for empty vector controls used in experiments with each strain. C) Spot densities for each strain on galactose (left) and glucose (right) media, normalized to the average spot density of empty vector controls used in experiments with each strain. Cross bars indicate mean ± standard error. The grey boundaries indicate the mean (solid line) and 95% confidence interval (grey rectangle) of spot densities for empty vector controls
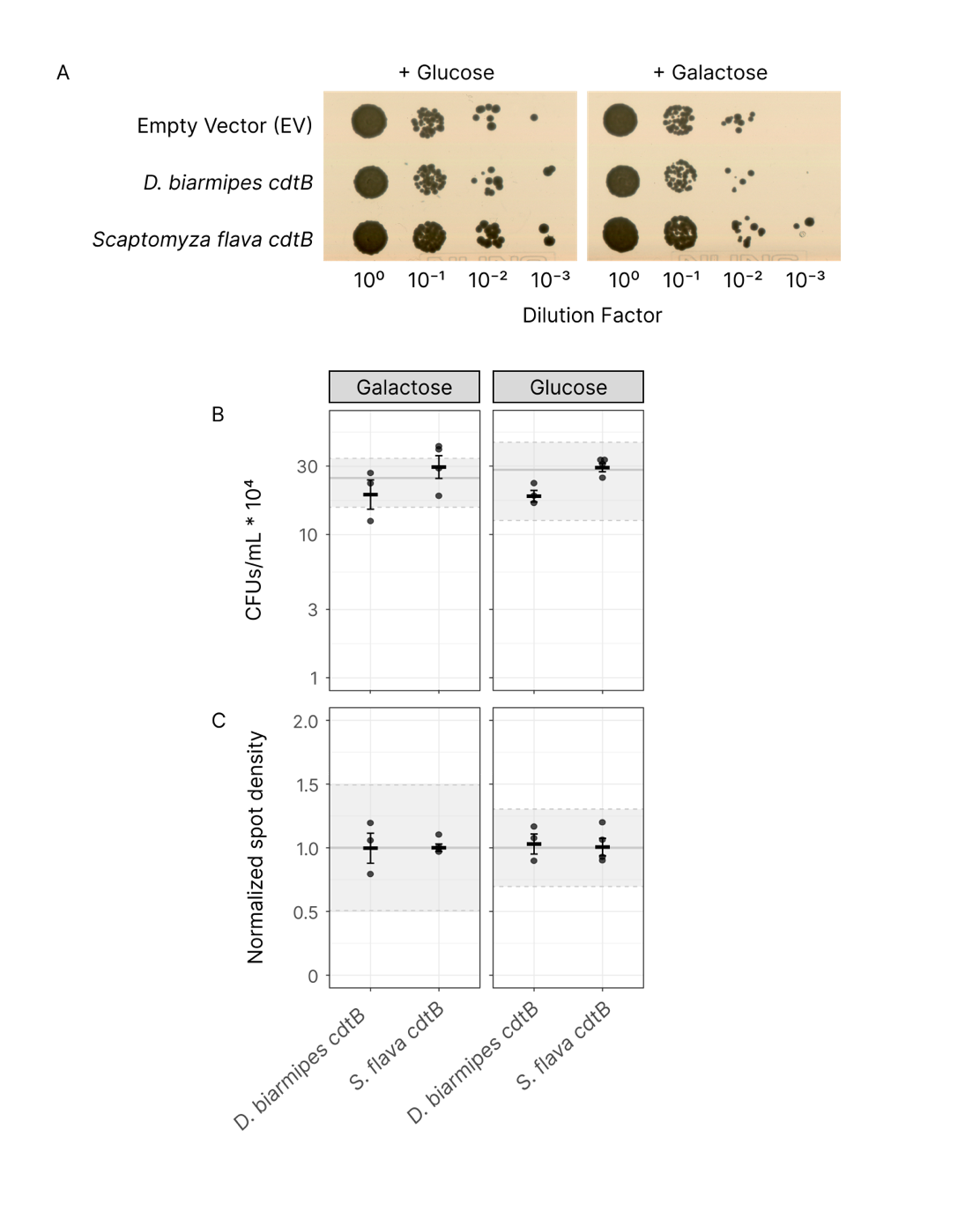

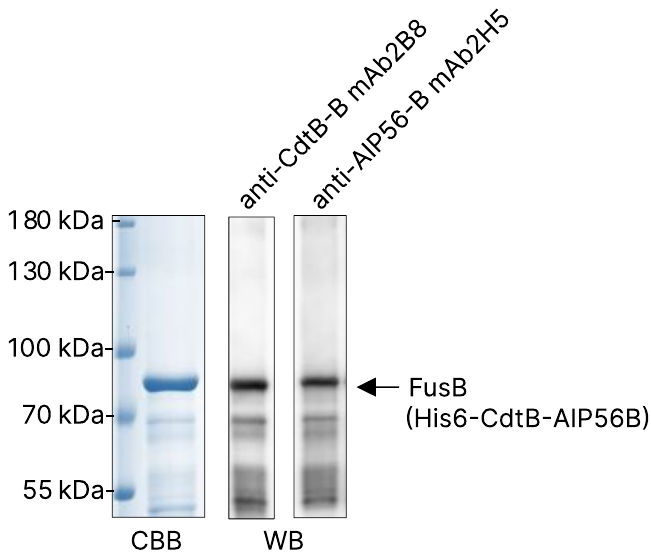
**Figure S16. SDS-PAGE analysis of the purified recombinant FusionB protein.** CBB= Coomassie brilliant blue. WB = western blot.

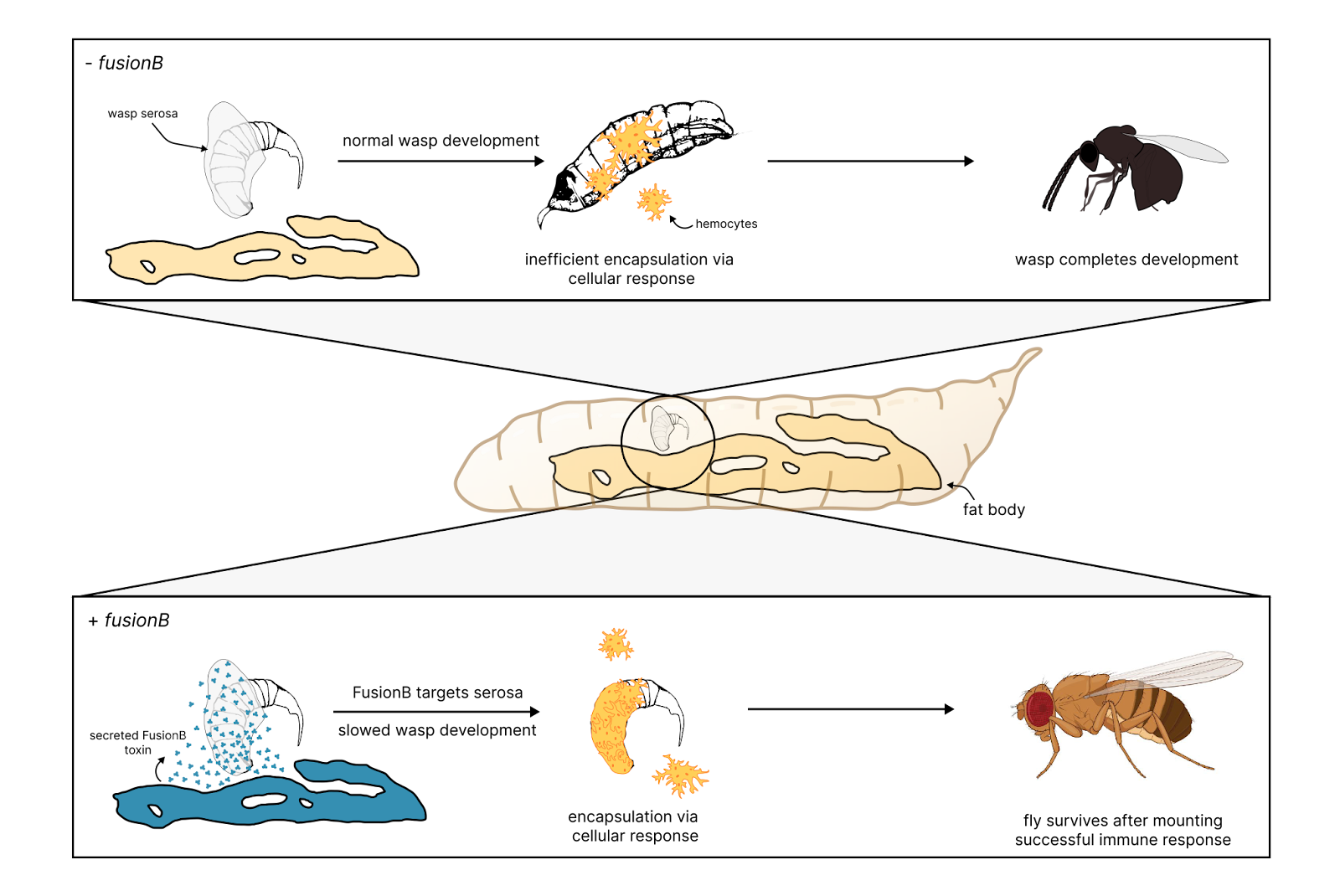
 **Figure S17. Model for FusionB involvement in parasitoid defense**. In the absence of FusionB, wasps can overcome the host immune response and complete development. When FusionB is secreted into the host hemolymph from the fat body, it binds to wasp serosal tissue and slows development of the immature wasp. Slowed wasp development buys time for other elements of the host response to act, allowing for more efficient and increased fly survival. Wasp neonate drawings are modified from Kopelman & Chabora (1984)^61^. This figure was created partially with BioRender.com.

| **Extended Data Table 1 – Parasitization Assay Statistics (per vial)** | | | | | | | | |
| --- | --- | --- | --- | --- | --- | --- | --- | --- |
|  |  | ***Uninfected controls*** | | | | | | |
|  |  | # vials | % larva to pupa | % larva to adult | % pupa to adult | % larva to wasp | % pupa to wasp | % both die |
|  | ***cdtB*** | 13 | 90.2 ± 5.6 | 89.3 ± 6.2 | 98.9 ± 1.3 | NA | NA | NA |
| **Cg-GAL4** | ***fusA*** | 10 | 92.9 ± 8.2 | 91.9 ± 8.0 | 99.0 ± 3.0 | NA | NA | NA |
|  | ***fusB*** | 19 | 90.3 ± 9.0 | 89.5 ± 8.9 | 99.1 ± 2.5 | NA | NA | NA |
|  | ***gfp*** | 14 | 85.2 ± 11.8 | 84.5 ± 11.8 | 99.1 ± 1.3 | NA | NA | NA |
|  |  | **Uninfected controls** | | | | | | |
|  |  | # vials | % larva to pupa | % larva to adult | % pupa to adult | % larva to wasp | % pupa to wasp | % both die |
| **He-GAL4** | ***fusB*** | 17 | 89 ± 14.3 | 87.8 ± 14.8 | 98.6 ± 2.7 | NA | NA | NA |
|  | ***gfp*** | 5 | 95 ± 4.9 | 93.7 ± 6.6 | 98.5 ± 2.5 | NA | NA | NA |
|  |  | ***L. boulardi*** | | | | | | |
|  |  | # vials | % larva to pupa | % larva to melanotic adult | % pupa to melanotic adult | % larva to wasp | % pupa to wasp | % both die |
|  | ***cdtB*** | 12 | 85.0 ± 7.1 | 2.8 ± 2.8 | 3.2 ± 3.2 | 39.0 ± 11.1 | 46.1 ± 16.6 | 56.1 ± 11.3 |
| **Cg-GAL4** | ***fusA*** | 14 | 88.8 ± 5.0 | 4.1 ± 3.7 | 3.9 ± 3.7 | 44.9 ± 13.8 | 52.4 ± 16.6 | 47.1 ± 11.6 |
|  | ***fusB*** | 20 | 90.0 ± 7.2 | 23.3 ± 10.0*** | 25.8 ± 10.7*** | 6.9 ± 5.6*** | 7.6 ± 5.6*** | 69.0 ± 10.2*** |
|  | ***gfp*** | 13 | 86.4 ± 9.9 | 3.7 ± 5.1 | 4.2 ± 5.6 | 45.1 ± 12.7 | 52.6 ± 14.4 | 47.2 ± 13.5 |
|  |  | ***L. boulardi*** | | | | | | |
|  |  | # vials | % larva to pupa | % larva to melanotic adult | % pupa to melanotic adult | % larva to wasp | % pupa to wasp | % both die |
| **He-GAL4** | ***fusB*** | 10 | 93.3 ± 5.5 | 10.5 ± 7.5* | 10.7 ± 7.5* | 12.6 ± 4.6 | 12.9 ± 4.7 | 76.9 ± 6.8 |
|  | ***gfp*** | 9 | 80.4 ± 19.0 | 3.9 ± 6.3 | 3.7 ± 5.8 | 14.7 ± 7.7 | 18.2 ± 9.4 | 81.4 ± 7.4 |
|  |  | ***L. heterotoma*** | | | | | | |
|  |  | # vials | % larva to pupa | % larva to melanotic adult | % pupa to melanotic adult | % larva to wasp | % pupa to wasp | % both die |
|  | ***cdtB*** | 10 | 91.2 ± 7.2 | 0 ± 0 | 0 ± 0 | 84 ± 7.0 | 92.2 ± 5.0 | 16 ± 7.0 |
| **Cg-GAL4** | ***fusA*** | 9 | 94.1 ± 6.4 | 0 ± 0 | 0 ± 0 | 89.2 ± 5.6 | 94.9 ± 3.9 | 10.8 ± 5.6 |
|  | ***fusB*** | 13 | 90.6 ± 6.0 | 0.6 ± 1.5 | 0.8 ± 1.8 | 57.0 ± 10.4*** | 63.0 ± 11.6*** | 42.2 ± 9.9*** |
|  | ***gfp*** | 16 | 86.7 ± 8.7 | 0.2 ± 0.6 | 0.2 ± 0.7 | 79.8 ± 9.9 | 92.0 ± 5.8 | 19.6 ± 9.8 |
|  |  | ***L. heterotoma*** | | | | | | |
|  |  | # vials | % larva to pupa | % larva to melanotic adult | % pupa to melanotic adult | % larva to wasp | % pupa to wasp | % both die |
| **He-GAL4** | ***fusB*** | 9 | 92.6 ± 4.6 | 0 ± 0 | 0 ± 0 | 59.4 ± 18.4* | 62.8 ± 18.1** | 40.6 ± 18.4* |
|  | ***gfp*** | 6 | 88.3 ± 11.6 | 0 ± 0 | 0 ± 0 | 76.7 ± 8.6 | 87.2 ± 6.3 | 23.3 ± 8.6 |
|  | * = p < 0.05, ** = p < 0.01, *** = p < 0.001 via pairwise Wilcoxin rank sum tests, relative to *gfp* controls. | | | | | | | |

**Table S1. Parasitization assay statistics for all experiments.** Mean proportion ± standard deviations are reported for all parasitization experiments.

| **Extended Data Table 2 — Parasitization Assay Raw Counts and Summary Statistics** | | | | | | | | | |
| --- | --- | --- | --- | --- | --- | --- | --- | --- | --- |
|  |  | **Uninfected controls** | | ***L. boulardi*** | | | ***L. heterotoma*** | | |
|  |  | # flies | # no emergence | # melanotic adult flies | # adult wasps | # no emergence | # melanotic adult flies | # adult wasps | # no emergence |
|  | ***cdtb*** | 691 | 7 | 29 | 338 | 452 | 0 | 504 | 96 |
| **Cg-GAL4** | ***fusa*** | 487 | 5 | 34 | 377 | 375 | 2 | 539 | 50 |
|  | ***fusb*** | 1016 | 10 | 280*** | 84*** | 809*** | 5 | 445*** | 328*** |
|  | ***gfp*** | 711 | 6 | 33 | 369 | 378 | 2 | 768 | 188 |
|  | **Chi-square (χ^2^)** | χ^2^ = 0.147, d.f. = 3, p = 0.986 | | **χ^2^ = 703.254, d.f. = 6, p < 2.2e-16** | | | **χ^2^ = 259.79, d.f. = 3, p < 2.2e-16** | | |
|  | **P-value  (Kruskal-Wallis test):** |  |  | **3.163E-08**  **( χ^2^ =37.769, d.f. = 3)** | **2.137E-08**  **( χ^2^ =38.573, d.f. = 3)** | **4.063E-06**  **( χ^2^ =27.768, d.f. = 3)** | 0.2115  ( χ^2^ =4.5088, d.f. = 3) | **1.655E-06  ( χ^2^ =29.717, d.f. = 3)** | **1.583E-06**  **( χ^2^ =29.717,**  **d.f. = 3)** |
|  |  | *** = p < 0.001 via pairwise Wilcoxin rank sum tests, relative to *gfp* controls. | | | | | | | |
|  |  | **Uninfected controls** | | ***L. boulardi*** | | | ***L. heterotoma*** | | |
| **He-GAL4** | ***fusb*** | 900 | 12 | 127 | 106 | 693 | 0 | 316 | 217 |
|  | ***gfp*** | 281 | 4 | 20 | 84 | 463 | 0 | 276 | 84 |
|  | **Chi-square (χ2)** | χ^2^ = 6.578e-28,     d.f. = 1, p = 1 | | **χ^2^ = 42.316, d.f. = 2, p = 6.474e-10** | | | **χ^2^ = 28.273, d.f. = 1, p = 1.054e-7** | | |
|  | **P-value  (Wilcoxin Test):** |  |  | **W = 73,**  **p = 0.023** | W = 38,  p = 0.595 | W = 30,  p = 0.234 | W = 27,  p = 1 | **W = 6.5,**  **p = 0.018** | **W = 47.5,**  **p = 0.018** |
|  |  | **Bold text** indicates tests with significant values | | | | | | | |

**Table S2. Parasitization assay raw count data and summary statistics.** Raw counts of flies, wasps, and “stalemate” pupae in each experiment.

| **Extended Data Table 3 – *Actin-Gal4* cross results** | | | | |
| --- | --- | --- | --- | --- |
| **Genotype** | **N (# crosses)** | **Observed proportion (%)** | **Expected proportion (%)** | **P-value (Exact binomial test)** |
| **; *Hd-cdtB/Actin-Gal4*** | 7 | 31.0 ± 6.3 | 33 | 0.1724 |
| **; *Da-cdtB/Actin-Gal4*** | 9 | 37.8 ± 4.6 | 33 | 0.9997 |
| **; *fusionA/Actin-Gal4*** | 6 | 38.7 ± 3.7 | 33 | 0.9989 |
| **; *fusionB/Actin-Gal4*** | 6 | 0.4 ± 0.7 | 33 | **< 2.2e-16** |
| **; +/*Actin-Gal4* ; *fusionA*/+** | 5 | 32.5 ± 7.3 | 25 | 1 |
| **; +/*Actin-Gal4* ; *fusionB*/+** | 4 | 0.4 ± 0.7 | 25 | **< 2.2e-16** |
|  | H_a_: observed < expected | | | |

**Table S3. *Actin-Gal4* cross results.** Observed and expected proportions of non-balancer offspring are reported. p-values reflect likelihood under the alternative hypothesis that fewer flies without the balancer phenotype were observed than expected (exact binomial test).
